# Distinct Chiral Nanostructures of Graphene Quantum Dots Govern Divergent Passive and Active Enantioselective Transport across Biological Membranes

**DOI:** 10.64898/2026.02.16.706189

**Authors:** Farbod Shirinichi, Yichen Liu, Runyao Zhu, James Carpenter, Wei Zhang, Yamil J. Colón, Yichun Wang

## Abstract

Chirality in two-dimensional nanomaterials provides a powerful lever to control biological interfaces, yet the structural origins of nanoscale chirality remain poorly understood. Here, we systematically investigate chiral ligand modulated graphene quantum dots (GQDs) and reveal how ligand stereochemistry and edge chemistry modulate the formation of diverse chiral or achiral nanostructures. Spectroscopy, microscopy, and density functional theory with ring-puckering analysis identified six structural motifs, twisted-, twisted-boat, saddle-shaped, hybrid, unbuckled, and random. Among these, twisted-, twisted-boat, and saddle-shaped GQDs exhibited genuine nanoscale structural chirality, while unbuckled, hybrid, and random conformations lacked organized distortion. Importantly, structural chirality governed passive permeation into biological membrane (e.g. lipid membrane of extracellular vesicles), whereas achiral variants relied mainly on hydrophobic interactions. In contrast, active transport across biological membrane (e.g. endocytosis) is insensitive to nanoscale structural chirality but strongly influenced by chiral ligand identity and transporter recognition. Collectively, these results establish chiral ligand conjugation as a modular route to program both chiral and achiral motifs in graphene nanostructures and highlight nanoscale structural chirality as a design principle for engineering bio-nano interactions.

## INTRODUCTION

Chirality, the property of non-superimposable mirror images, is a defining feature of living systems. Majority of the biomolecules in nature, including proteins, nucleic acids, sugars, and lipids are chiral, and this handedness governs molecular recognition, catalysis, and information transfer across hierarchical length scales.^1,2^ Biological membranes likewise exhibit chiral organization, as asymmetric lipid packing and protein crowding generate interfacial energy landscapes that favor one enantiomeric interaction over the other.^3,4^ At the nanoscale, this chiral order of lipid membrane couples to curvature, tilt, and lateral-pressure profiles, leading to helicity-dependent adsorption, orientation, and barrier-crossing pathways for approaching objects.^4,5^ Given that biomolecular assemblies and biological membranes are inherently nanoscale, exploiting structural chirality of nanomaterials offers a versatile strategy for regulating their transmembrane transport, including passive permeation, and active tissue uptake.^6–8^ These considerations highlight the importance of treating nanomaterial chirality not simply as a descriptor but as a controllable design variable for biomedical applications.^9^

Carbon nanostructures have been widely studied for their chiral conformations and for leveraging distinct interactions at biointerfaces across diverse biomedical applications.^10–12^ Their chirality can arise either from chiral precursors used to construct the nanocarbon or from post-synthetic functionalization with chiral ligands on carbon nanostructures.^11,13,14^ Graphene quantum dots (GQDs), quasi-zero-dimensional carbon nanostructures with excellent photoluminescence, photostability, and biocompatibility, and abundant surface sites for functionalization,^15–17^ have been investigated extensively and demonstrated to possess tunable chiral properties.^8,18^ While majority of these effects originate from the molecular chirality of surface functional groups or conjugated ligands, local edge chemistry can impose strain fields that distort the basal plane of GQDs and thereby induce structural chirality. Early demonstrations attached chiral ligands such as cysteine to GQD edges, producing chiral GQDs with a twisted nanostructure which exhibit strong circular dichroism signals.^18,19^ Such nanoscale chirality has been demonstrated to facilitate transport across the tumor extracellular matrix (ECM),^6,7^ and improve the cargo loading and delivery efficiencies of drug-loaded small extracellular vesicles (sEVs).^8,16,20^ Despite significant progress, the various types of ligand-encoded distortions that can be induced, and the extent to which they can be precisely controlled, remain unclear due to the vast diversity and distinct physicochemical properties of available chiral ligands.^21^ Hence, the range of accessible chiral and achiral geometries of GQDs and their consequent impact on bio-nano interactions, remain insufficiently characterized. How these distinct chiral nanostructures beyond surface chemistry govern passive and active enantioselective transport across biological membranes remains to be elucidated.

Herein, we investigated a library of chiral ligands, including 18 amino acids with diverse properties, which induced ligand stereochemistry into six well-defined nanostructures of GQDs (**Figure 1a**), thereby imparting distinct biointerfacial behaviors. By combining chiroptical spectroscopy and microscopy with Density Functional Theory (DFT) and molecular dynamics (MD) simulations, together with Cremer-Pople ring puckering analysis,^22^ we resolve six characteristic GQD nanostructures: twisted, twisted boat, saddle shaped, hybrid, unbuckled, and disordered (**Figure 1b**). Studies of ligand-modulated chiral GQD interacting with sEV lipid bilayers demonstrate that structural chirality governs passive membrane permeation, with different chiral conformation and distortion degrees yielding distinct transmembrane efficiencies, while achiral structures exhibit non-enantioselective entry via nonspecific hydrophobic insertion at relatively low efficiencies (**Figure 1c**). In contrast, cell uptake of chiral GQDs primarily via endocytosis is insensitive to chiral structures and instead reflects the biochemical identity of the conjugated amino acids, consistent with the transporter recognition. Overall, these findings uncover the vast varieties of chiral nanostructures accessible in GQDs and establish structural chirality, rather than surface chemistry alone, as a governing parameter for passive transmembrane transport. This work expands the structural landscape of carbon nanomaterials and provides design principles for selecting edge chemistries and distortion amplitudes to tune passive and active enantioselective transport across biological membranes. These insights suggest practical routes toward enantioselective probes, chiral drug carriers, and imaging platforms that exploit the inherent chirality of biological membranes.

**Figure 1.**
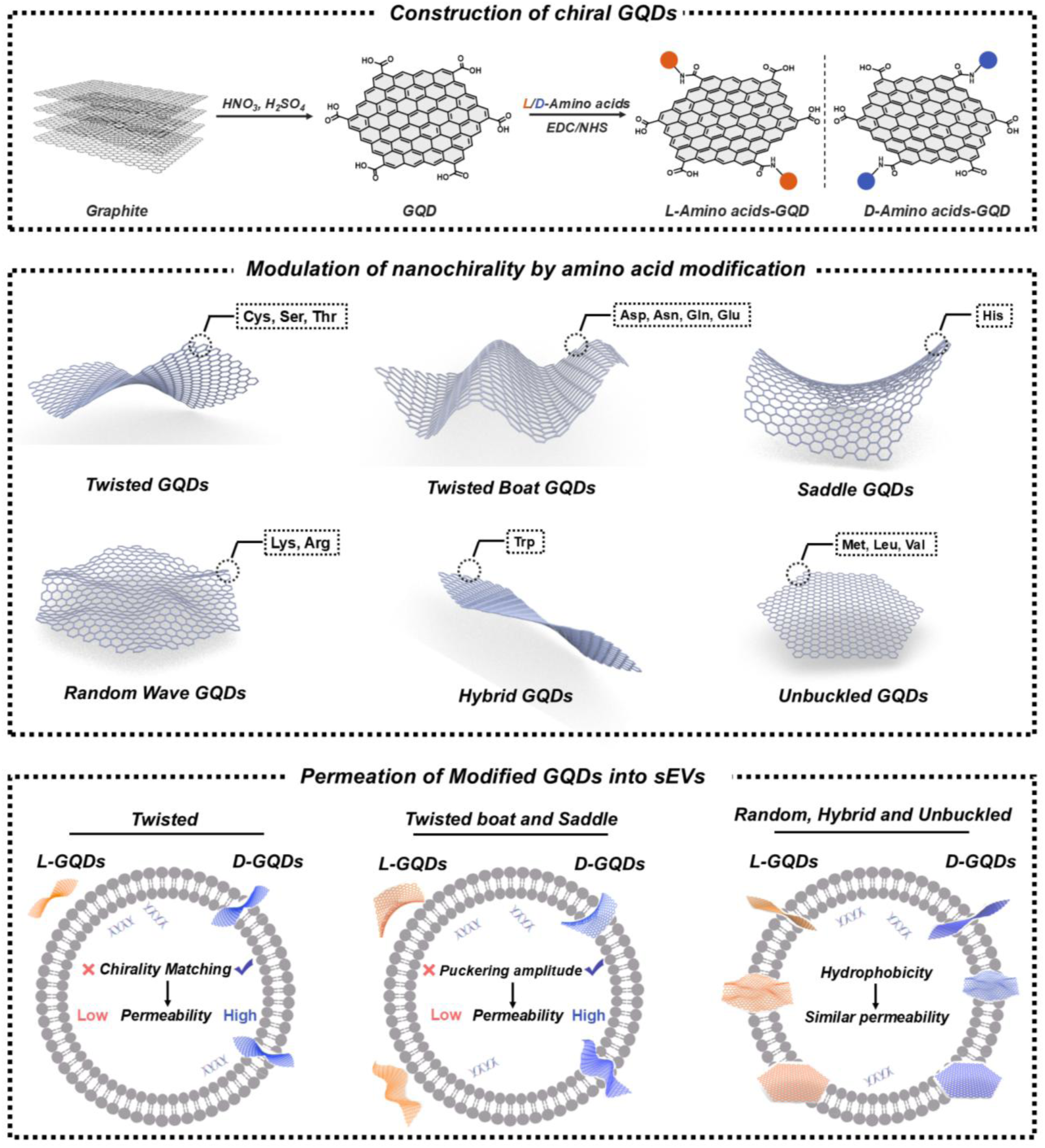
a) Synthesis route of graphene quantum dots (GQDs) via oxidative cutting of carbon nanofibers, followed by chiral ligand conjugation to form chiral GQDs. b) Classification of induced chiral nanostructures based on ligand types; scheme not to scale and exaggerated for clarity. c) Comparison of permeability efficiency into lipid membrane of small extracellular vesicles (sEVs) among different GQD structures, highlighting differences in passive transmembrane transport between *L-* and *D-*amino acid conjugates.

## RESULTS AND DISCUSSION

### Synthesis and Characterization of Chiral Ligand-Modulated GQDs

To investigate how ligand chirality is transferred to nanomaterials and thereby influences its structures, chiroptical properties, and biological interactions, we systematically functionalized GQDs with a comprehensive library of chiral ligands (amino acids). For 14 amino acids, Cys, Ser, Thr, His, Asp, Asn, Glu, Gln, Trp, Met, Val, Leu, Arg, and Lys, both enantiomers were successfully conjugated to GQDs, yielding distinct chiroptical activities. GQDs were synthesized via oxidative cutting of carbon nanofibers following our previously reported procedure^23^, resulting in uniformly dispersed nanosheets. Transmission electron microscopy (TEM) confirmed that the as-synthesized GQDs possess an average lateral dimension of approximately 6.2 nm and exhibit a well-defined graphene crystalline structure, as evidenced by visible lattice fringes with an interplanar spacing of 0.242 nm (**Figure 2a**).^24^ Edge functionalization was achieved by covalently attaching *L-* and *D-*ligands through 1-Ethyl-3-(3-dimethylaminopropyl)carbodiimide (EDC) and N-Hydroxysuccinimide (NHS) chemistry.^8,18^ Fourier-Transform Infrared Spectroscopy (FTIR) analysis confirmed successful conjugation (**Figure 2b; Figure S1-3**). In all chiral ligand functionalized GQDs, new absorption bands appeared at ∼1730, ∼1545 and ∼1250 cm⁻¹, corresponding to amide bond vibrations^25–27^. Additional peaks between 2900–3050 cm⁻¹ were attributed to aliphatic C–H stretching^27^, further indicating incorporation of amino acid side chains. For Gln- and Asn-functionalized GQDs, overlapping intrinsic amide signals complicated interpretation^28^, yet distinct spectral features at fingerprint region confirm the covalent linkage to the GQD framework. Atomic force microscopy (AFM) and TEM were used to examine the morphological changes of ligand functionalized GQDs. While TEM measurements showed consistent lateral sizes after conjugation (∼6.4 nm across variants; **Figure 2c**; **Figure S4-6**), AFM revealed differences in vertical thickness. Most variants remained similar to pristine GQDs with average height of 1.17nm,^24^ but His, Cys, Ser, Trp-functionalized GQDs displayed substantially greater heights increase in range of 0.51–0.83 nm, (**Figure 2d; Figure S7-9; Table S1**), suggesting out-of-plane deformations or clustering at edges. Zeta potential measurements showed that the edge functionalization of GQD reduced the density of free carboxyl groups and may affect the negative surface charges.^18,20^ Positively charged ligands (Arg, Lys, His) shifted the potential toward positive values, whereas negatively charged ones (Asp, Glu) offset the loss of carboxyl groups, resulting in minimal net change (**Figure 2f**). Other amino acid conjugates showed intermediate decreases of negative charges, consistent with edge conjugation. Taking together, our results confirm that amino acids covalently bind to GQD edges through amide formation, tuning interfacial chemistry and surface charge while preserving the graphene core. AFM reveals localized out-of-plane distortions, especially in His-, Cys-, and Ser-functionalized GQDs, suggesting the onset of structural chirality.

The optical properties of chiral ligand functionalized GQDs were examined to evaluate how edge functionalization perturbs the electronic environment. Photoluminescence (PL) spectra revealed red*-*shifted emission of all functionalized samples compared to that of pristine GQDs, with maxima in the 450–550 nm range when excited at 360 nm (**Figure 2e** and **Figure S10**). This shift is attributed to increased edge defects and reduced band gap energy following conjugation, consistent with trends observed in other π-conjugated nanomaterials where chemical modification alters defect states and electronic transitions.^18,29,30^ UV–vis absorption spectra provided complementary evidence. All functionalized GQDs exhibited a new band at ∼255 nm, absent in pristine samples (**Figure 2g** and **Figure S11**).^31^ This feature originates from perturbation of the electronic states of the graphene backbone by the conjugated chiral ligands that corresponds to the π–π* transitions of the sp^2^ hybridized aromatic backbone,^18,32^ with the red*-*shifted emission previously attributed to hybridized orbitals at the edges and the possible formation of exciplex-like states in aromatic domains of the GQDs.^18,30,33^ Together, optical spectroscopy confirms that ligand conjugation modifies the local electronic structure at GQD edges, as evidenced by red-shifted PL and new UV–vis absorption bands arising from orbital hybridization, edge defects, and exciton confinement, demonstrating tunable optical properties through ligand chemistry. Circular Dichroism (CD) spectroscopy was used to study the emergent chirality in the functionalized GQDs (**Figure 2g** and **Figure S12-15**). While strong signals between 190–225 nm corresponded primarily to the molecular chirality of ligands^8,34,35^, additional shifts and inversions reflected structural changes in the GQD nanostructure.^18,32,35^ Three categories of spectral behavior were observed among all samples, reflecting differences in ligand properties and stereochemistry. The positively charged amino acids, Arg- and Lys-conjugated GQDs exhibited broadened, red-shifted D bands due to electrostatic interactions, steric repulsion, and hydrogen bonding around the amide bonds.^21,36–39^ Certain polar amino acids, such as Asp, Asn, Gln, and Glu, also produced a red shifted and narrower CD band, suggesting hydrogen-bonding effects (**Figure S12**).^21^ Hydrophobic amino acids, including Trp, Leu, Met, and Val, induced secondary CD peaks with the same sign as the chiral ligands that diminished over time, potentially attributed to additional π–π stacking or van der Waals interactions.^37–41^ Notably, several special amino acids, including Cys-, His-, Thr-, and Ser, induced secondary peaks with the opposite sign to those of the free chiral ligands, indicating significant conformational rearrangements induced by edge conjugation.^18,37,41^

**Figure 2.**
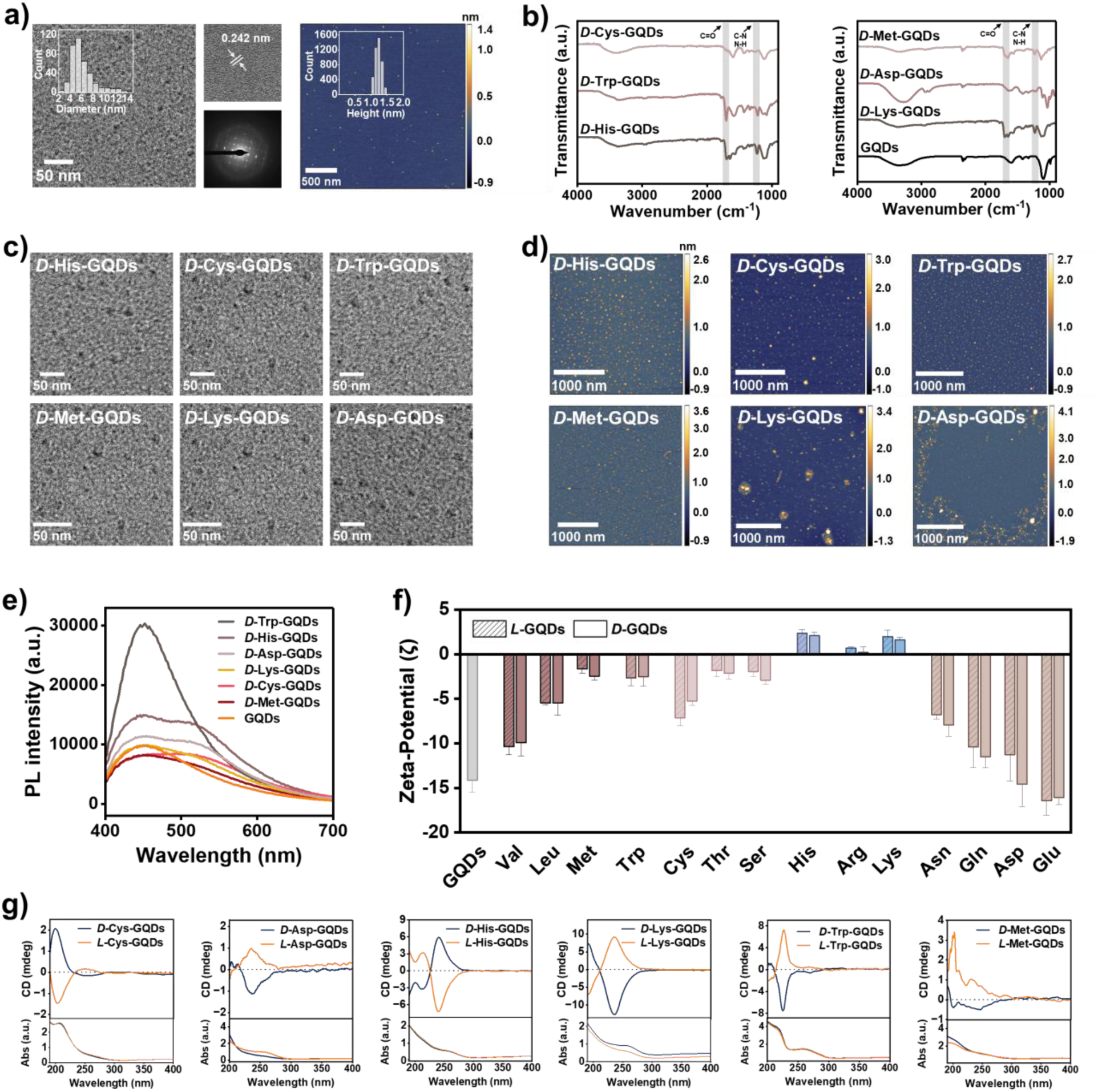
a) Transmission electron microscopy (TEM) and atomic force microscopy (AFM) image of synthesized GQDs. b) Fourier-Transform Infrared Spectroscopy (FTIR) spectra of pristine and amino acid*-*functionalized GQDs. c) TEM images of selected GQDs. d) AFM images of the same GQDs. e) Photoluminescence spectra of pristine and functionalized GQDs. f) Zeta potential comparison of chiral GQDs. g) Experimental CD spectra of representative amino acid–functionalized GQDs showing distinct spectral features associated with ligand identity and stereochemistry.

### Mechanistic Classification of Chiral*-*Ligand-Modulated GQD Nanostructures

CD spectra indicated three dominant chiroptical response types, but it could not resolve the underlying structural heterogeneity underlying these signals. We therefore combined DFT (including TD-DFT) with quantitative ring-puckering analysis to identify the nanoscale conformations responsible for the observed CD features. ^42,43^ Geometry optimization revealed six recurring GQD motifs: twisted, saddle-shaped, twisted-boat, hybrid, unbuckled, and random (**Figure 3a** and **Figure S16**). TD-DFT reproduced the experimental CD/UV–vis spectral envelopes, including peak inversions and relative intensities, supporting these structural assignments despite a systematic underestimation of absolute wavelength (**Figure 3b** and **Figure S17**).^18,44^ To quantify out-of-plane distortion and handedness, optimized structures were analyzed using ring puckering parameters: the total puckering amplitude (***Q***) reports deviation from planarity, and the phase angle (***φ***) encoded the handedness of distortion (**Figure 3f; see Supplemental Methods; Table S2-3**).^45,46^ Across all ligands, stable chiral nanostructures were obtained for Cys, Ser, Thr, His, Asp, Asn, Glu, and Gln, whereas Arg, Lys, Trp, Met, Leu, and Val predominantly yielded disordered or weakly distorted conformations that relaxed toward effectively achiral states. Trp-GQDs were a notable transient case: folding driven by π–π stacking between the indole ring and the GQD surface produced short-lived structural chirality, consistent with MD simulations indicating strong van der Waals adsorptions (**Figure 3g, h**).

**Figure 3.**
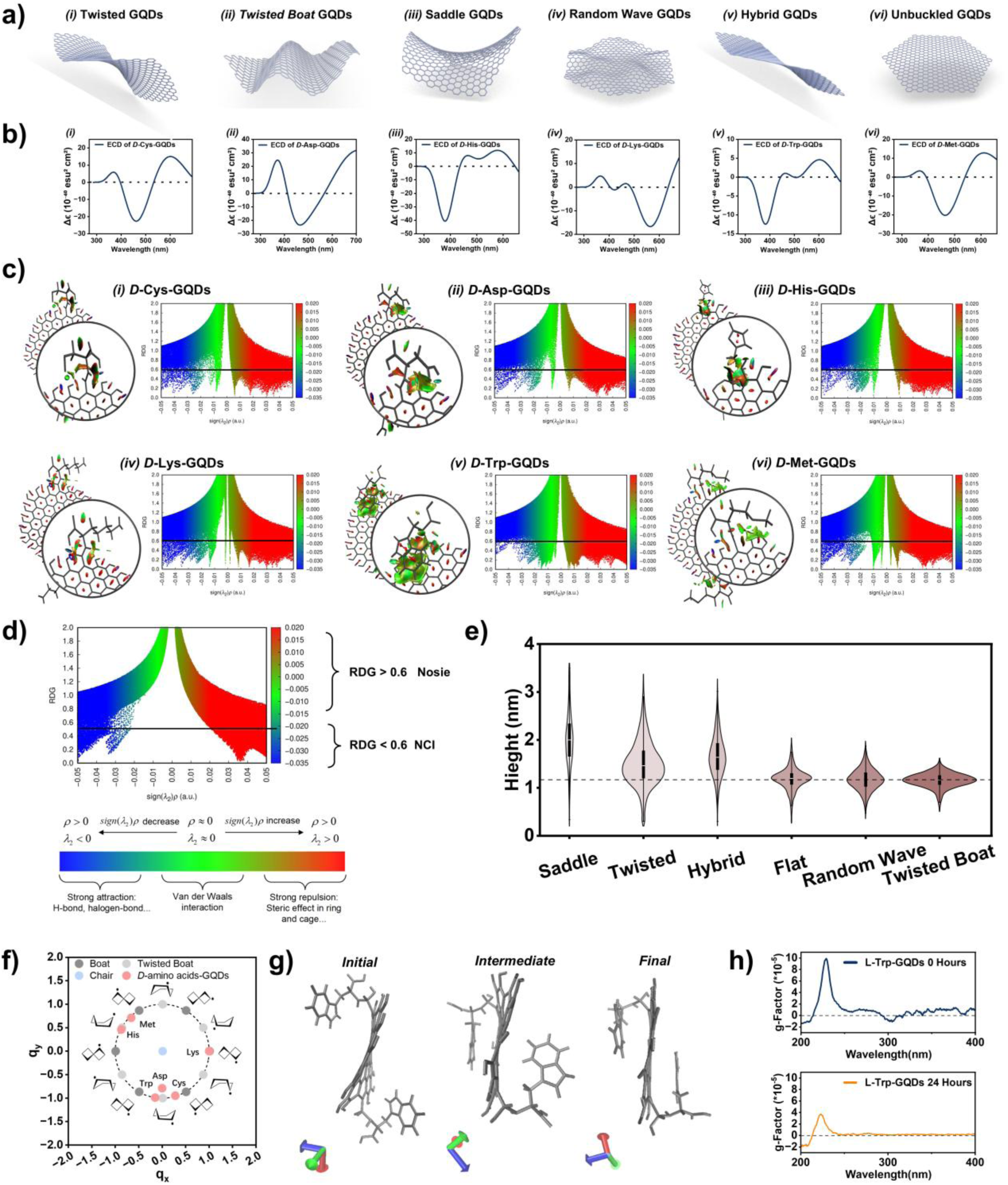
Structural classification of chiral ligand modulated GQDs by Circular Dichroism (CD) spectroscopy, Density Functional Theory (DFT) modeling, and ring puckering analysis. a) Geometries of selected GQDs illustrating structural motifs, including twisted-, twisted-boat, saddle-shaped, hybrid, random, and unbuckled conformations. scheme not to scale and exaggerated for clarity b) NCI/RDG isosurface mapping of six GQD motifs (twisted, twisted-boat, saddle-shaped, hybrid, random, and unbuckled), revealing distinct noncovalent interaction patterns associated with the formation of each structural motif. c) NCI/RDG analysis of unmodified GQDs, showing that noncovalent interactions are dominated by edge hydrogen bonding, while steric repulsion between benzene rings suppresses out-of-plane distortion. d) Violin plot showing AFM-derived height distribution across different chiral nanostructure groups e) Simulated electronic circular dichroism (ECD) spectra obtained from TD-DFT calculations, demonstrating good agreement with experimental CD data f) Puckering coordinate map (***qₓ***, ***qᵧ***) of representative *D*-amino acid–functionalized GQDs, illustrating their distribution into twisted-, twisted-boat, and saddle conformations. e) Molecular dynamics (MD) simulations of *D*-Trp-GQDs showing folding of the hydrophobic side chain over the GQD surface, which contributes to the observed dihedral distortion and enhanced chiral optical response h Time-dependent CD spectra of L-Trp-GQDs recorded at two different time points, showing gradual relaxation of the chiral structure, consistent with the conformational evolution observed in MD simulations, ultimately leading to loss of chirality.

Integrating experimental CD spectroscopy, AFM morphology, and DFT modeling with ring-puckering analysis shows that chiral ligand (amino acid) functionalized GQDs reproducibly populate these six conformational classes (**Figure 3a** and **Figure S16**). Three motifs (twisted, twisted-boat, and saddle-shaped) exhibit persistent structural chirality, whereas the remaining three (random, hybrid, and unbuckled) display transient or effectively achiral distortions. Because all systems share the same covalent amino-acid linkage, we attribute these divergent outcomes primarily to side-chain–backbone noncovalent interactions. Reduced Density Gradient (RDG) and NCI analysis further indicate that the spatial organization and directionality of these interactions, rather than their mere presence, determine propagates coherently into a stable, handed GQD nanostructure (**Figure 3c**).

*Stable chiral motifs.* **Twisted GQDs** arise primarily from small polar ligands (Cys, Ser, Thr), where local buckling at the attachment region propagates through the π-framework to generate a global twist (**Figure 3a-i** and **Figure S16-i**). Cys- and Ser-GQDs exhibit the largest puckering amplitudes (***Q*** = 0.6–0.9; **Table S2**) and increased heights in AFM relative to pristine GQDs (**Figure 3e** and **Table S1**), consistent with substantial out-of-plane distortion. Their CD spectra show an amino-acid band near ∼216 nm and a secondary inverted band at 250–285 nm assigned to the twisted GQD backbone (**Figure 2g** and **Figure S14**). RDG/NCI map of Cys-GQD indicates cooperative stabilization of overall structure by attractive contributions (e.g. multiple peaks at blue region) with distinct van der Waals interactions (e.g. 2 peaks at green region), and modest steric peak at sign(*λ*_2_)*ρ*=0.005 which shaped the twisted structure (**Figure 3b-i**). Thr-GQDs retain structural chirality by CD and DFT but show weaker puckering (***Q*** = 0.41; **Table S2**) and less height change, suggesting a smaller but still chiroptically effective deformation. **Twisted-boat GQDs** form mainly by ligands with polar/negatively charged side chains (Asp, Asn, Gln, Glu; **Figure S12**). These chiral ligands favor dense bond that led to preferable directional bending of ligands to the carboxylic edge of GQDs (**Figure 3b-ii**). RDG/NCI map shows single van der Waals peak at sign(*λ*_2_)*ρ*=-0.0075, localized hydrogen-bonding at sign(*λ*_2_)*ρ*=-0.025, and steric force signature with a single peak at sign(*λ*_2_)*ρ*=0.03 (**Figure 3b-ii**), consistent with modest puckering (***Q*** ≈ 0.4; **Table S2**), red-shifted CD bands without an inverted secondary peak (**Figure S12**), and negligible height changes (**Table S1**). His-GQDs uniquely produces a **saddle-shaped** topology driven by the combined effects of buckling strain and steric/electrostatic influence from the positively charged imidazole side chain (**Figure 3b-iii**). DFT predicts large distortions (***Q*** > 1.0; **Table S2**), consistent with characteristic CD features and increased heights (**Figure 3e** and **Figure S14** and **Table S1**). RDG/NCI maps show comparatively weaker attraction forces and limited van der Waals continuity but strong, continuous steric contributions, favoring an open saddle deformation rather than a compact twist (**Figure 3b-iii** and **Figure S16-iii**).

*Transient/achiral motifs.* Positively charged Arg and Lys produce **random, rippled GQDs** lacking persistent handedness (**Figure 3b-iv**). Their broadened, red-shifted CD signals (*g*-factors of 10⁻⁴; **Figure S13**) are consistent with localized distortions near amide linkages rather than coherent lattice chirality; RDG/NCI map shows intense, but spatially multiple random interaction peaks feature that impede propagation into an organized global deformation (**Figure 3b-iv**). Trp forms a **hybrid GQD** conformation via π–π stacking between the indole ring and the GQD π-system (**Figure 3b-v**), generating transient folding and a short-lived three-band CD signature (∼305, 270, 220 nm; **Figure 3g, h** and **Figure S15**) that decays within 24 h alongside only slight height increases (**Figure 3e**). Consistent with this behavior, DFT and MD indicate progressive relaxation towards flatten conformations driven by van der Waals-dominated stabilization and hydrophobic collapse (**Figure 3b-v, g**). Hydrophobic ligands (Met, Leu, Val) yield an **unbuckled GQD** conformation, near-planar state with only minor edge perturbations (**Figure 3a-vi, b-vi**, **Figure S15** and **Figure S16-iv**), weak time-dependent CD signals near 210–265 nm, and no measurable height changes (**Table S1**). RDG/NCI mapping features are diffuse and lack the bending of amino acids toward the edge of GQDs that are required for stable out-of-plane deformation (**Figure 3a-vi, b-vi, Figure S16-vi, Figure S18** and **Table S3**).

Enantioselective coupling between ligand configuration and GQD handedness was observed only when stable chiral motifs formed (Cys, Ser, Thr, His, Asp, Asn, Glu, Gln): *L*-enantiomers produced right-handed distortions and *D*-enantiomers yielded left-handed ones (**Figure 2g**). In systems that do not stabilize coherent lattice deformation, optical activity primarily reflects the intrinsic molecular chirality of the ligands rather than GQD-level structural chirality. Racemic conjugation eliminates the distortions (**Figure S19**), confirming stereochemistry as the essential driver. Collectively, these results establish that six ligand-dependent GQD conformational classes governed by side-chain–backbone noncovalent interaction balance. Twisted, twisted-boat, and saddle-shaped motifs maintain persistent structural chirality through coherent, directional interaction networks, whereas random, hybrid, and unbuckled motifs fail to be stabilized sustained out-of-plane deformation and therefore exhibit weak or transient chiroptical responses.

### Distinct Passive Transmembrane Transport of Chiral*-*Ligand Modulated GQDs

Biological membranes provide the largest interfacial area for molecular exchange and functional regulation in living systems. Their intrinsic enantioselectivity gives rise to distinct transmembrane behaviors for *L*- and *D*-enantiomers through both passive and active pathways. Understanding transmembrane transport is therefore essential for the rational design of nanomaterials intended to interface with biological systems. To elucidate how structural chirality governs membrane interactions, we first investigated the passive permeation of chiral-ligand–modulated GQDs through lipid bilayers, using sEVs as a biologically relevant model that captures lipid packing features but lacks cellular energy-dependent transport machinery (**Figure 4a-b** and **Figure S20-22**).^8,47^ The resulting permeation efficiencies revealed two mechanistic regimes: one dominated by chirality-driven structural interactions and the other by hydrophobicity-driven insertion ^48,49^

For GQDs with pronounced stable structural chirality (twisted, twisted-boat, and saddle-shaped), permeation was strongly influenced by chirality matching with the lipid bilayer. Because biological membranes exhibit intrinsic left-handed chirality, left-handed GQDs (*φ* < 0) (**Table S3**), generated by conjugation with *D*-amino acids (e.g., *D*-Cys, *D*-Ser, *D*-Thr, *D*-Asp, *D*-Asn, *D*-Glu, *D*-Gln, *D*-His), showed higher permeation than their enantiomeric counterparts.^3,18,48,50,51^ The extent of this enhancement depended on the structural origin of chirality. In twisted GQDs (Cys-, Ser-, Thr-functionalized), puckering originates in the GQD core, inducing a continuous helical distortion across the basal plane (**Figure 3a-i** and **Figure S16-i**).^18^ Within this class, permeation increased with puckering amplitude (***Q***), where larger ***Q*** promoted better chiral alignment and more efficient permeation via improved insertion (**Figure 4e; Table S3**).^8^ In contrast, twisted-boat and saddle-shaped GQDs derive chirality primarily from edge deformations (**Figure 4f**), leading to distinct geometrical constraints and permeation behaviors. Twisted-boat GQDs (Asp, Asn, Gln, Glu) showed among the highest permeation efficiencies (60-75%) despite their relatively mild chiral distortions (*Q* < 0.5), suggesting an optimal balance between chiral asymmetry and membrane packing compatibility (**Figure 4f**).^52^ By comparison, saddle-shaped *D*-His-GQDs exhibited reduced permeation (29–48%). Their larger vertical distortions (Q > 1.0) create rigid protrusions, consistent with an AFM height of ∼2 nm (**Figures S7–S9; Table S1**), which hinders passive permeation. This subtle asymmetry appears to provide an optimal balance between structural chirality and compatibility with membrane packing.^50,52^

For GQDs lacking distinguishable structural chirality (e.g., Met-, Leu-, Val-, Arg-, Lys-, Trp-GQDs), lipid membrane permeation was governed primarily by hydrophobic interactions.^53,54^ Variants such as Met-, Leu-, and Val*-*GQDs exhibited higher permeation efficiencies through sEVs membranes than Arg- and Lys-GQDs, correlating positively with their hydrophobicity (**Figure 4c**).^53,55^ Interestingly, Trp-GQDs, despite their high hydrophobicity, showed the lowest permeation. This is primarily due to the unique anchoring behavior of Trp side chains at the bilayer interface, where they stabilize via interfacial polarization rather than insertion, thereby hindering complete translocation. Consistent with this mechanism, Trp-GQDs remained associated with membrane surfaces but were readily detached upon washing, resulting in low measured permeation.^56,57^ Further, we used trypan blue, a membrane-impermeant fluorescence quencher, to probe the location of chiral Trp-GQDs on sEVs. After normalization to the respective unquenched controls (set to 100%), only ∼30% of the initial signal remains for both L-Trp-GQDs at sEVs and D-Trp-GQDs at sEVs (**Figure S23**). This pronounced loss indicates that chiral Trp-GQDs are predominantly surface-associated rather than inserted into the lipid bilayer, consistent with the low permeation observed in independent assays.

**Figure 4.**
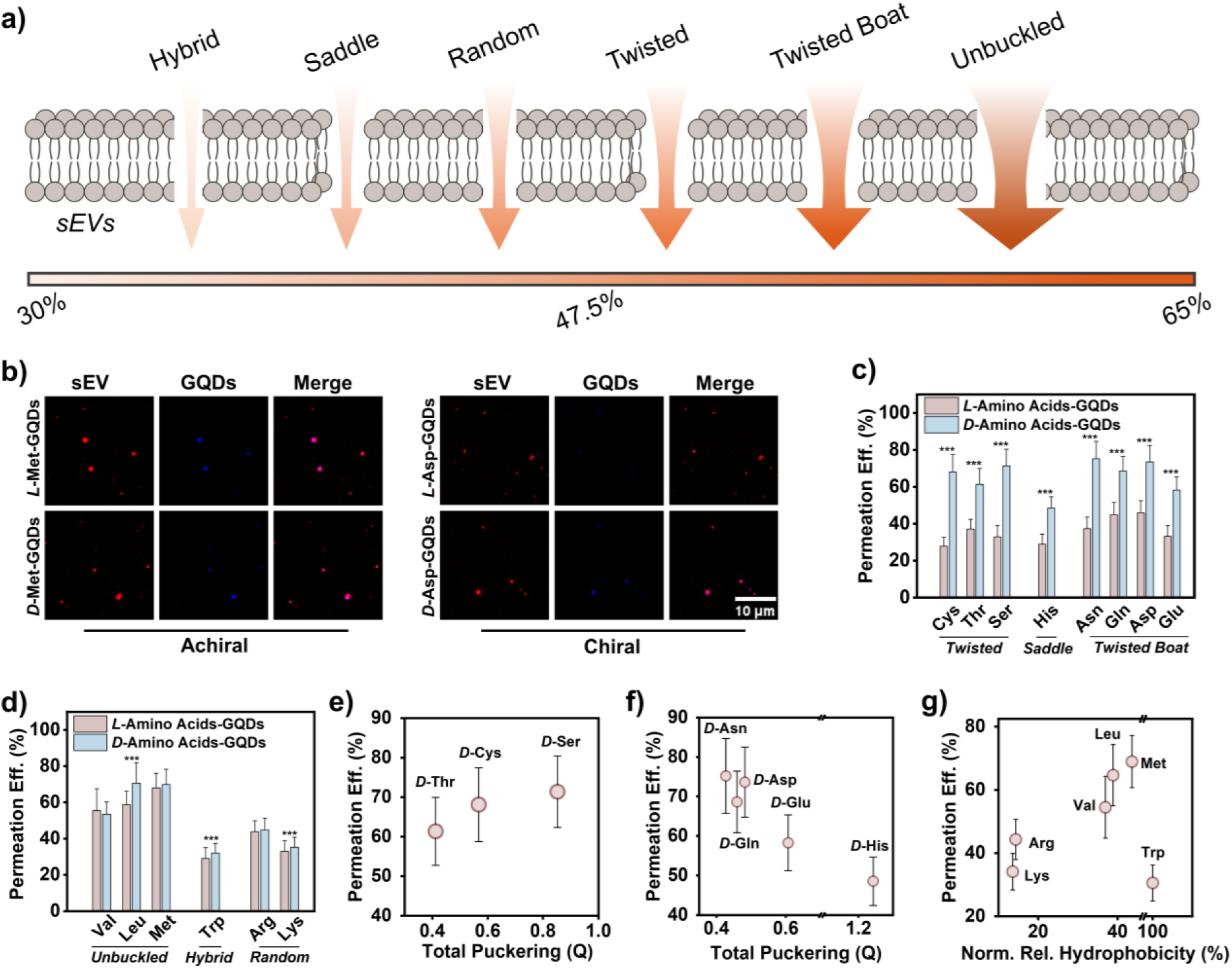
Permeation of chiral ligand functionalized GQDs into sEVs. a) Schematic illustration of GQD interaction with the lipid bilayer, highlighting different permeation of nanostructural classes. b) Representative confocal images of sEVs incubated with selected chiral and achiral ligand functionalized GQDs, showing differences in internal fluorescence intensity. DiD labels the plasma membrane. Scale bars: 10 μm. Quantification of permeation efficiency for c) chiral nanostructures (twisted-, saddle, twisted-boat) and d) achiral nanostructures (unbuckled, hybrid, random). (*n* = 5). Correlation between permeation efficiency and puckering amplitude (Q) for e) twisted-boat, saddle-shaped GQDs, f) twisted-GQDs, and g) achiral GQDs. (*n* = 5). Data are mean ± SD. Statistical differences were analyzed by a Student’s two-sided t-test. \*\*\**P*<0.001.

Together, these findings reveal that nanoscale chirality governs passive permeation when ordered nanostructures are present, while hydrophobicity dominates insertion for disordered or achiral systems. Twisted- and twisted-boat GQDs exhibit favorable chirality matching with lipid bilayers, while saddle-shaped GQDs experience geometry-induced mismatches that hinder diffusion. Moreover, differences between twisted-, twisted-boat, and saddle-shaped nanostructures highlight the importance of the spatial origin of chirality in nanostructures. These insights establish chirality as a powerful design principle for tuning passive transmembrane transport, with broad implications for nanomedicine.

### Transporter-Dominant Cell Uptake of Chiral*-*Ligand Modulated GQDs

Cell membranes contain numerous active transport pathways, raising the possibility that chiral nanostructures may have influence transmembrane transport of living cells differently. To examine this, we quantified cellular uptake of previously characterized chiral ligand modulated GQDs in both HepG2 (cancerous) and THLE-2 (healthy) cell lines. For most GQD systems, HepG2 cells showed higher uptake of *L*-GQDs than *D*-GQDs, whereas THLE-2 cells showed uniformly minimal differences between *L* and *D* (**Figure 5**). This pattern contrasts with the enantioselective permeation observed across sEV membranes, where passive transport is strongly governed by chirality matching, puckering amplitude, and hydrophobicity. To determine whether the HepG2 enantioselective bias depends on active internalization, we repeated the assay at 4 °C, which suppresses ATP-dependent endocytosis and largely restricts interactions to membrane association at the cell surface. Under this condition, total cellular fluorescence decreased in both cell types and the *L*–*D* differences collapsed to non-significant (**Figure S24**). These results indicate that the chirality dependence observed in HepG2 at 37 °C arises from energy-dependent internalization, rather than passive partitioning into the bilayer. In living cells, uptake of GQDs is therefore dominated by protein-mediated internalization, and ligand–protein recognition together with ligand stereochemistry can outweigh nanoscale structural handedness.^58^

**Figure 5.**
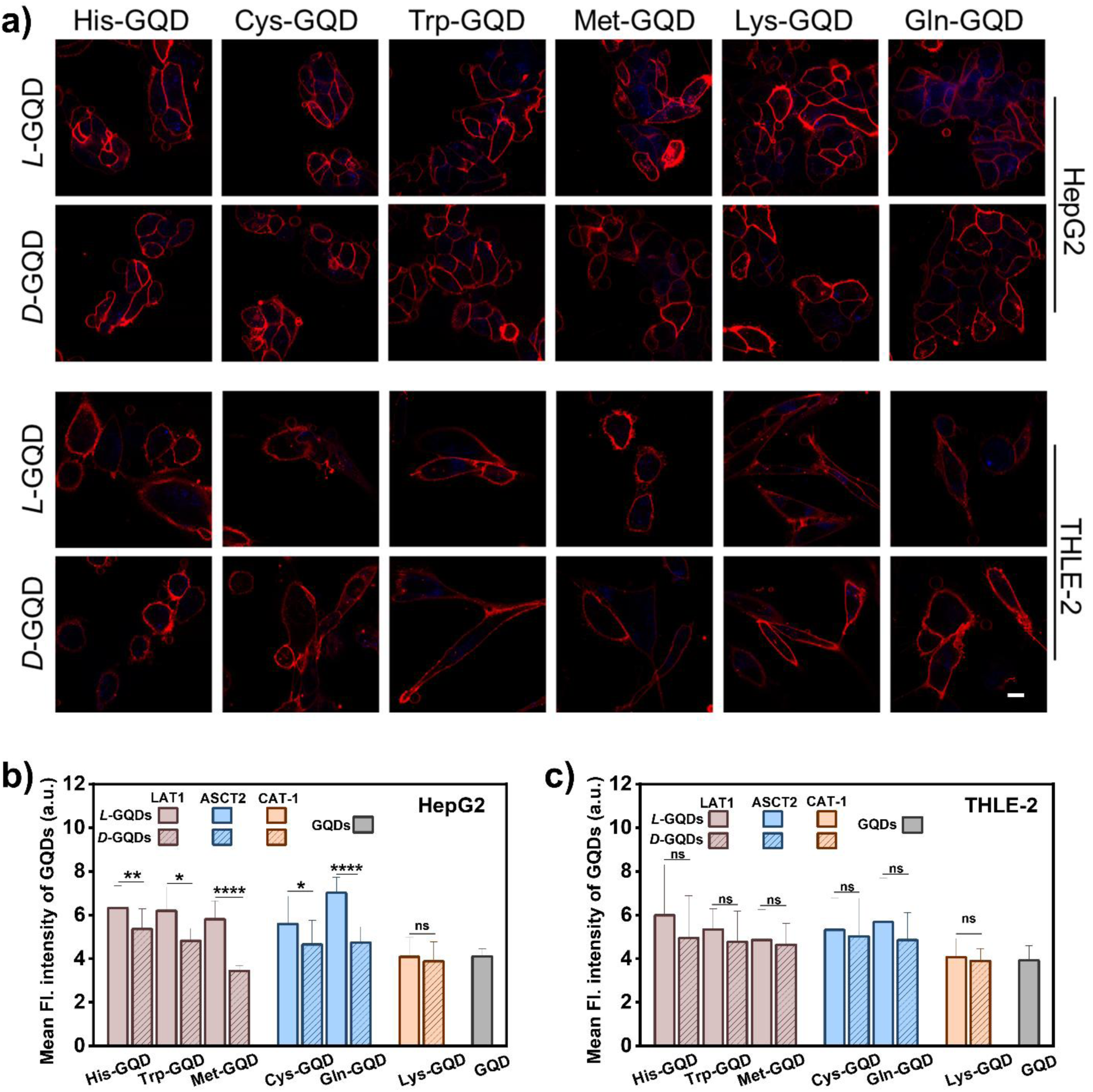
Cellular uptake of amino-acid–conjugated GQDs at 37 ℃. a) Representative confocal images of HepG2 and THLE-2 cells after 4 h incubation with the indicated chiral GQD variants (5 μM). DiD labels the plasma membrane. Scale bars: 10 μm. (*n* = 5). Quantification of cellular GQD fluorescence for b) HepG2 and c) THLE-2 corresponding to panel a). Data are mean ± SD. Statistical differences were analyzed by a Student’s two-sided t-test. \**P*<0.05, \*\**P*<0.01, \*\*\**P*<0.001.

For active transmembrane transport, the chirality of surface chemical motif plays a more dominant role than the nanoscale structural conformations, driving selective endocytosis in transporter-rich cells while limiting uptake in healthy counterparts.^59,60^ This clear distinction between passive, bilayer-mediated permeation and active, transporter-mediated uptake through biological membrane offers unique opportunities to selectively target lipid-enveloped viruses or vesicles without inducing cytotoxicity in host cells. For example, because the lipid envelopes of many viruses resemble sEV membranes in their packing characteristics and lack active transport machinery, twisted or mildly twisted *D*-GQDs could be inserted into the viral membrane to deliver antiviral agents or disrupt membrane integrity, while minimizing entry into healthy cells. In parallel, exploiting transporter-guided tumor targeting and chiroptical imaging or diagnostic readouts provides complementary strategies for achieving high selectivity. Overall, these findings highlight the potential of integrating nanoscale chirality and molecular stereochemistry to design multifunctional nanomaterials for selective therapeutic delivery and pathogen inhibition.

## CONCLUSION

Chiral ligand conjugation programs GQDs into six distinct nanostructures, among which twisted-, twisted- boat, and saddle geometries exhibit genuine structural chirality at nanoscale, while hybrid, random, and unbuckled variants reflect transient or disordered distortions, the results are highly reproducible across independent syntheses and repeated characterizations. Stereochemistry dictated handedness, with *L-* and *D-*forms producing mirror-image conformations and racemic mixtures abolishing organized distortions. These structural classes directly influenced interactions with lipid bilayers. Chiral nanostructures permeated membranes enantioselectivity through chirality matching, whereas achiral variants relied primarily on hydrophobic insertion. This distinction establishes structural chirality at nanoscale as a governing principle for passive and selective permeation across lipid bilayers and for modulating their interactions with graphene quantum dots. Beyond elucidating the origin of nanoscale chirality in GQDs, this work establishes generalizable rules linking nanostructure geometry to biological transport, clarifying how specific structural distortions govern membrane permeation and enantioselectivity.

## AUTHOR INFORMATION

### Corresponding Author

*Corresponding author, Professor Yichun Wang

Department of Chemical & Biomolecular Engineering University of Notre Dame Notre Dame, IN 46556, USA

### Author Contributions

Y.W. supervised the entire project, including overall conceptualization, validation, data analysis, and manuscript preparation. F.S. designed the methodology and performed all experiments, including the synthesis and characterization of GQDs and chiral ligand–modulated GQDs, as well as all analytical chemistry work, data analysis, and manuscript writing. Y.L. designed and integrated the computational component, carried out the DFT calculations, and contributed to manuscript writing. Y.L. and R.Z. performed the biological and drug-loading experiments, comparing the chiral ligand–modulated GQD method for sEVs loading and conducting cell-uptake studies. R.Z. isolated and characterized sEVs and carried out confocal imaging. J.C. and Y.C. performed the MD simulations for the chiral ligand–modulated GQDs. W.Z. designed and tested the attachment of Trp-GQDs to the sEVs membranes.

Farbod Shirinichi^+^ and Yichen Liu^+^ contributed equally

## ACKNOWLEDGMENT

This work was financially supported by funding from the National Science Foundation. The authors gratefully acknowledge the facilities at the University of Notre Dame, including the Analytical Science and Engineering at Notre Dame (ASEND), the Center for Research Computing (CRC), the Electron Microscopy Core, the Optical Microscopy Core, and the Center for Environmental Science and Technology (CEST), for providing instrumentation and assistance with characterization and imaging throughout this research.

## ABBREVIATIONS

AA: Amino Acids
Arg: Arginine
Asp: Aspartic acid
Asn: Asparagine
Cys: Cysteine
Glu: Glutamic acid
Gln: Glutamine
His: Histidine
Leu: Leucine
Lys: Lysine
Met: Methionine
Ser: Serine
Thr: Threonine
Trp: Tryptophan
Val: Valine

