## Supporting Information for "Distinct Chiral Nanostructures of Graphene Quantum Dots Govern Divergent Passive and Active Enantioselective Transport across Biological Membranes"

### Materials and Methods

#### Chemicals and Reagents

Sulfuric acid ($H_2SO_4$, 98%), nitric acid ($HNO_3$, 68%), sodium hydroxide (NaOH), and carbon nanofibers were obtained from Sigma-Aldrich and used as received without further purification. Deionized (DI) water was used in all aqueous preparations. All experiments were conducted under ambient conditions unless otherwise noted.

#### Synthesis of Graphene Quantum Dots (GQDs)

Graphene quantum dots were synthesized via oxidative acid treatment of carbon nanofibers following a previously reported protocol with modifications. Briefly, 0.45 g of carbon nanofibers were added to 90 mL of concentrated $H_2SO_4$ (98%) in a 250 mL two-neck roun*D*-bottom flask and stirred for 1.5 h. Subsequently, 30 mL of concentrated $HNO_3$ (68%) was added dropwise, maintaining an $H_2SO_4$: $HNO_3$ volume ratio of 3:1. The resulting mixture was sonicated for 1 h in an ice bath, ensuring the temperature remained below 35 °C.

The oxidized dispersion was then transferred to a reflux apparatus and heated at 120 °C for 21–24 h. The completion of the reaction was indicated by a color change from black to golden yellow. After cooling to room temperature, the mixture was neutralized to pH 6.5–7.0 by the gradual addition of 5 M NaOH (~700 mL). The neutralized solution was filtered through a 0.22 μm membrane and dialyzed against DI water using a 2000 Da MWCO membrane for 3 days with water changes every 8 h. The resulting GQD dispersion was stored at 4 °C for further use.

**Synthesis of Chiral Amino Acid–Modified GQDs**

Chiral GQDs were synthesized via EDC/NHS-mediated covalent conjugation of various L- and *D*-amino acids to hydroxyl-functionalized GQDs (H-GQDs). The amino acids used included L-/*D*-Arg, His, Lys, Asp, Glu, Gln, Asn, Ser, Thr, Cys, Val, Ile, Leu, Met, and Trp. Reaction times varied between 24 and 72 h depending on the amino acid.

For conjugation, H-GQDs (0.5 mg/mL) were activated by adding 0.8 µL of 100 mM EDC, followed by stirring for 30 min at 0°C. NHS (36.2 µL, 500 mM) was subsequently added, and the mixture was stirred for an additional 30 min to form the active ester. A 100 mM aqueous solution of the amino acid (36.2 µL) was then added, and the reaction mixture was stirred at room temperature for 24–72 h. To enhance dispersion, brief sonication (5 min) was performed every 6 h during the reaction. The crude product was purified using centrifugal filtration (2 kDa MWCO, 6000 rpm, 30 min, 3 cycles) to remove unreacted ligands and byproducts. Conjugation was confirmed by FTIR and CD spectroscopy.

The amino acids used included L-/*D*- Ser, Val, Ile, Leu, Met, and Trp. Reaction times varied between 24 and 72 h depending on the amino acid. For conjugation, H-GQDs (0.5 mg/mL) were activated by adding 0.8 µL of 100 mM EDC, followed by stirring for 30 min at room temperature. NHS (36.2 µL, 500 mM) was subsequently added, and the mixture was stirred for an additional 30 min to form the active ester. A 100 mM aqueous solution of the amino acid (36.2 µL) was then added, and the reaction mixture was stirred at room temperature for 24–72 h. To enhance dispersion, brief sonication (5 min) was performed every 6 h during the reaction. The crude product was purified using centrifugal filtration (2 kDa MWCO, 6000 rpm, 30 min, 3 cycles) to remove unreacted ligands and byproducts. Conjugation was confirmed by FTIR and CD spectroscopy.

The amino acids used included L-/*D*- Asn, and Glu. Reaction times varied between 72 h depending on the amino acid. For conjugation, H-GQDs (0.5 mg/mL) were activated by adding 0.8 μL of 100 mM EDC, followed by stirring for 30 min at 0°C and at pH 10. NHS (36.2 μL, 500 mM) was subsequently added, and the mixture was stirred for an additional 30 min to form the active ester. A 100 mM aqueous solution of the amino acid (36.2 μL) was then added, and the reaction mixture was stirred at room temperature for 24–72 h. To enhance dispersion, brief sonication (5 min) was performed every 6 h during the reaction. The crude product was purified using centrifugal filtration (2 kDa MWCO, 6000 rpm, 30 min, 3 cycles) to remove unreacted ligands and byproducts. Conjugation was confirmed by FTIR and CD spectroscopy.

**Transmission Electron Microscopy (TEM)**

TEM characterization was performed using a Thermo Scientific™ Talos F200i microscope operated at 200 kV. For sample preparation, 5 μL of a 30 μM dispersion of pristine or amino-acid-modified GQDs were dropped-cast onto carbon-coated 300-mesh copper TEM grids. The grids were left undisturbed for 10 minutes to allow adsorption of the GQDs, the excess droplet was gently wicked off using filter paper, and the samples were then followed by air-drying for 1 hour at room temperature. Subsequently, the grids were placed in a vacuum desiccator and kept under reduced pressure for 24–72 hours to ensure complete solvent removal and prevent surface contamination prior to imaging.

**Atomic Force Microscopy (AFM)**

Surface morphology and height distribution of pristine and amino-Acid-modified GQDs were examined using a Bruker Jupiter XR atomic force microscope operated in tapping mode. Freshly cleaved mica substrates were used for sample deposition. A 10 μL aliquot of a 30 μM dispersion of GQDs or modified GQDs was drop-cast onto the mica surface and left undisturbed for 10 minutes to allow adsorption. The excess droplet was gently wicked off using filter paper, and the samples were then air-dried for 8–24 hours at room temperature before imaging. AFM measurements were performed using an AC55 cantilever tip with a scan resolution of 512 × 512 pixels. Image processing and quantitative analyses were carried out using Gwyddion software, from which height profiles, surface roughness, and topographical distributions were extracted.

**UV–Vis Absorption and Photoluminescence (PL)**

UV–Vis absorption spectra were recorded using a JASCO J-1700 circular dichroism spectrometer equipped with a 0.1 cm quartz cuvette. All measurements were performed in deionized (DI) water containing mixtures of pristine or amino-acid-modified GQDs at concentrations ranging from 0.5 to 5 μM, with a total sample volume of 200 μL. Photoluminescence (PL) spectra were obtained using a SpectraMax i3x multi-mode microplate reader at GQD concentrations of 0.5 to 15 μM, using an excitation wavelength of 365 nm and collecting emission spectra from 430 to 700 nm. Samples were loaded into black-bottom 96-well plates with a total volume of 200 μL per well. All spectra were recorded at room temperature and were baseline-corrected and normalized to account for background and instrumental response.

**Circular Dichroism (CD)**

CD spectra were recorded using a JASCO J-1700 circular dichroism spectrometer equipped with a 10 mm (0.1 cm) quartz cuvette. Measurements were performed in deionized (DI) water containing mixtures of pristine and amino-acid-modified GQDs at concentrations ranging from 0.5 to 5 μM, with a total sample volume of 200 μL. Spectra were acquired at room temperature over the 190–400 nm range using an appropriate bandwidth and scan rate to ensure stable signal acquisition. Baseline correction was carried out using DI water as the reference, and spectra were normalized to the same optical density for comparison with UV–Vis absorption measurements.

**Fourier Transform Infrared (FTIR) Spectroscopy**

FTIR spectra were acquired using a Shimadzu IRXross spectrometer. For GQD and amino-acid-modified GQD samples, 2 µL of a 50 µM aqueous dispersion was drop-cast directly onto the FTIR crystal and left to air-dry for 30 minutes at room temperature prior to measurement. Spectra were then collected after complete evaporation of the droplet to ensure film formation on the crystal surface. For comparison, pure amino acid samples were analyzed in their solid powder form under identical instrumental conditions. All spectra were recorded in the mid-infrared range with appropriate background subtraction and baseline correction applied before analysis.

**Zeta Potential**

Zeta potential measurements were performed using a Malvern Zetasizer Nano ZS instrument. Samples consisted of 5 µM aqueous dispersions of pristine or amino-acid-modified GQDs prepared in deionized (DI) water. Measurements were conducted at room temperature, and each sample was analyzed in triplicate to ensure reproducibility. The Hückel approximation was applied to calculate the electrophoretic mobility and corresponding zeta potential values, with instrumental parameters adjusted to account for the size and surface characteristics of the GQDs.

**Density Functional Theory (DFT) and Electronic Circular Dichroism (ECD) Calculations**

Geometry optimizations were performed using the B3LYP functional. TD-DFT calculations were employed to simulate ECD spectra. Experimental CD spectra were used to guide iterative refinement of the initial GQD–amino acid geometries. Only structures with consistent spectral features between computed and measured ECD were used for final structural assignments.

Puckering parameters (Q, $q_x$, $q_\gamma$, φ) were calculated to assess nanostructural deformation and chirality.

#### sEVs Permeation Assay

To evaluate the permeation efficiency of chiral GQDs into small extracellular vesicles (sEVs), 150 µL of 3T3-sEVs (~$1 \times 10^9$ particles/mL) and 50 µL of chiral GQDs were mixed to a final GQD concentration of 15 µM. The mixture was incubated at room temperature for 20 min to allow interaction and potential uptake.

Following incubation, the samples were washed three times with PBS (8 °C) using a 100 kDa centrifugal filter to remove unincorporated free GQDs. For imaging, 5 µL of the sEV suspension was mounted between 18 mm × 18 mm coverslips and imaged using confocal laser scanning microscopy (CLSM) under the DAPI fluorescence channel (excitation/emission: 405/425–525 nm) with a 100× oil-immersion objective. Z-stack imaging was performed across four randomly selected fields of view, with each Z-stack consisting of 30 optical slices collected at 0.125 µm intervals.

The total number of fluorescent sEV particles (TFEPs) was quantified using manually adjusted fluorescence thresholds, calibrated to match the observed size distribution of sEVs. Quantification was performed using the Batch Process mode of the imaging software, using a standardized code with the corresponding threshold settings applied uniformly across all samples.

#### Trypan blue quenching assay

3T3-sEVs (150 µL; ~$1 \times 10^9$ particles/mL) were mixed with chiral Trp-GQDs (50 µL; final15 µM) and incubated at room temperature for 20 min. Samples were then washed three times with PBS (8 °C) using a 100 kDa centrifugal filter to remove unincorporated GQDs. The resulting L-Trp-

GQDs@EVs and D-Trp-GQDs@EVs were incubated with trypan blue solution (0.4%, 40μL) for 5 min in microcentrifuge tubes. The fluorescence of each sample was measured and expressed as a percentage relative to the corresponding unquenched control.

**Ring Puckering formulation**

The puckering coordinates describe out-of-plane deformations of a local six-membered carbon ring on the GQD edge or basal plane, adapted from Cremer–Pople definitions.

**i:** Index of the six carbon atoms forming a local six-membered ring unit on the edge or basal plane of the GQD.

**$z_i$:** Out-of-plane displacement (Å) of the *i*th carbon atom relative to the mean plane defined by the six-membered ring; positive values indicate upward buckling, negative downward values.

**Mean Plane:** Defined by least-squares fitting of the six ring carbon atoms; serves as the reference plane for measuring puckering distortions.

**$q_2 \cdot \sin(\varphi)$:** In-plane puckering component calculated as

$$q_2 \cdot \sin(\varphi) = \frac{1}{\sqrt{3}} \times \sum_{i=1}^{6} z_i \cdot \sin\left(\frac{2\pi(i-1)}{3}\right)$$

**$q_2 \cdot \cos(\varphi)$:** In-plane puckering component orthogonal to $q_2 \cdot \sin(\varphi)$, given by

$$q_2 \cdot \cos(\varphi) = \frac{1}{\sqrt{3}} \times \sum_{i=1}^{6} z_i \cdot \cos\left(\frac{2\pi(i-1)}{3}\right)$$

**$q_2$:** Magnitude of the second order puckering component representing in-plane deformation intensity,

$$q_2 = \sqrt{(q_2 \cdot \sin(\varphi))^2 + (q_2 \cdot \cos(\varphi))^2}$$

**$q_3$:** Third order puckering coordinate describing out-of-plane buckling of the six-membered carbon ring,

$$q_3 = \frac{1}{\sqrt{6}} \times \sum_{i=1}^{6} z_i \cdot (-1)^{i-1}$$

**Q (Total Puckering Amplitude):** Total puckering amplitude representing the overall curvature of the local GQD region,

$$Q = \sqrt{q_2^2 + q_3^2}$$

**$q_x$, $q_\gamma$ (Puckering Direction Components):** Normalized directional components of the puckering vector within the GQD plane,

$$q_x = \frac{q_2 \cdot \sin(\varphi)}{Q}$$

$$q_y = \frac{q_2 \cdot \cos(\varphi)}{Q}$$

**φ (Phase Angle):** Phase angle describing the nature of the deformation pattern and handiness of structure

$$\mathbf{Tan}(\varphi) = \frac{q_\gamma}{q_x}$$

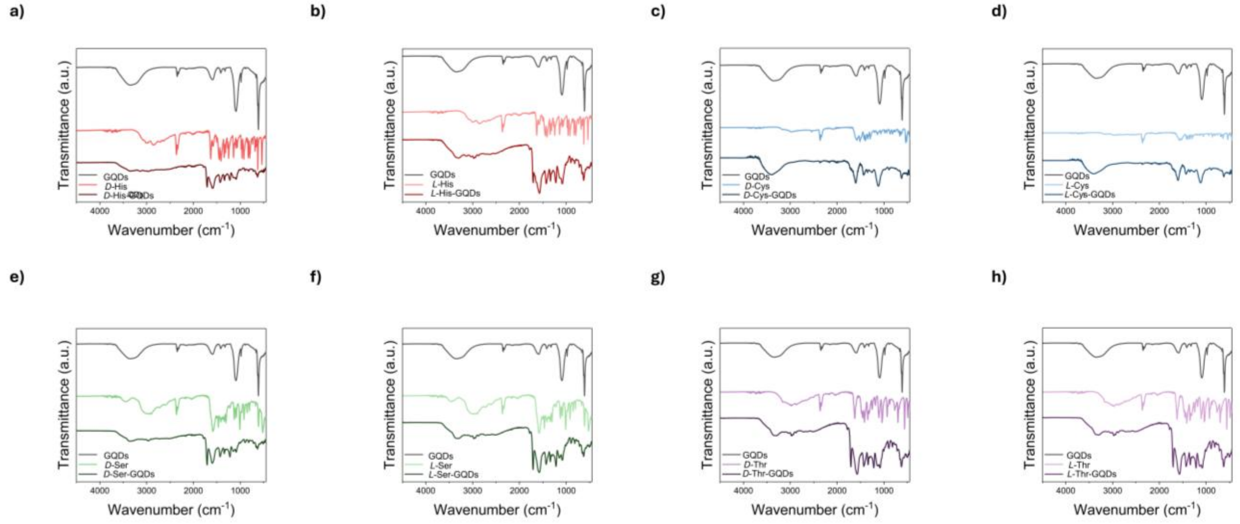


**Figure S1.** Fourier Transform Infrared (FTIR) spectra of pristine graphene quantum dots (GQDs) and *D*- or *L*-amino acid–modified GQDs. a) *D*-His–GQDs, b) *L*-His–GQDs, c) *D*-Cys–GQDs, d) *L*-Cys–GQDs, e) *D*-Ser–GQDs, f) *L*-Ser–GQDs, g) *D*-Thr–GQDs, and h) *L*-Thr–GQDs. Characteristic amide vibrations at ~1730, 1545, and 1250 $cm^{-1}$ and C–H stretching bands at 2900–3050 $cm^{-1}$ confirm covalent conjugation of amino acids to GQD edges.

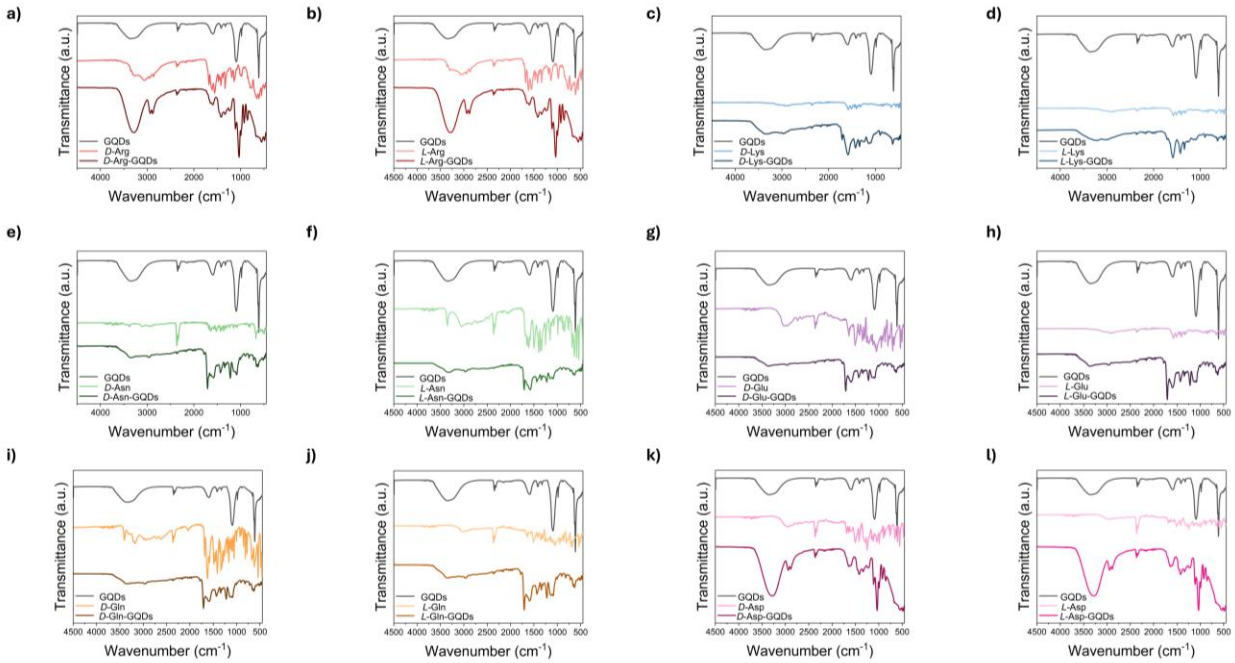


**Figure S2.** FTIR spectra of pristine GQDs and *D*- or *L*-amino acid–modified GQDs. a) *D*-Arg–GQDs, b) *L*-Arg–GQDs, c) *D*-Lys–GQDs, d) *L*-Lys–GQDs, e) *D*-Asn–GQDs, f) *L*-Asn–GQDs, g) *D*-Glu–GQDs, h) *L*-Glu–GQDs, i) *D*-Gln–GQDs, j) *L*-Gln–GQDs, k) *D*-Asp–GQDs, and l) *L*-Asp–GQDs. All modified samples exhibit new absorption bands at ~1730, 1545, and 1250 $cm^{-1}$ corresponding to amide bonds, along with peaks at 2900–3050 $cm^{-1}$ from C–H stretching, confirming successful conjugation of amino acids to GQD edges.

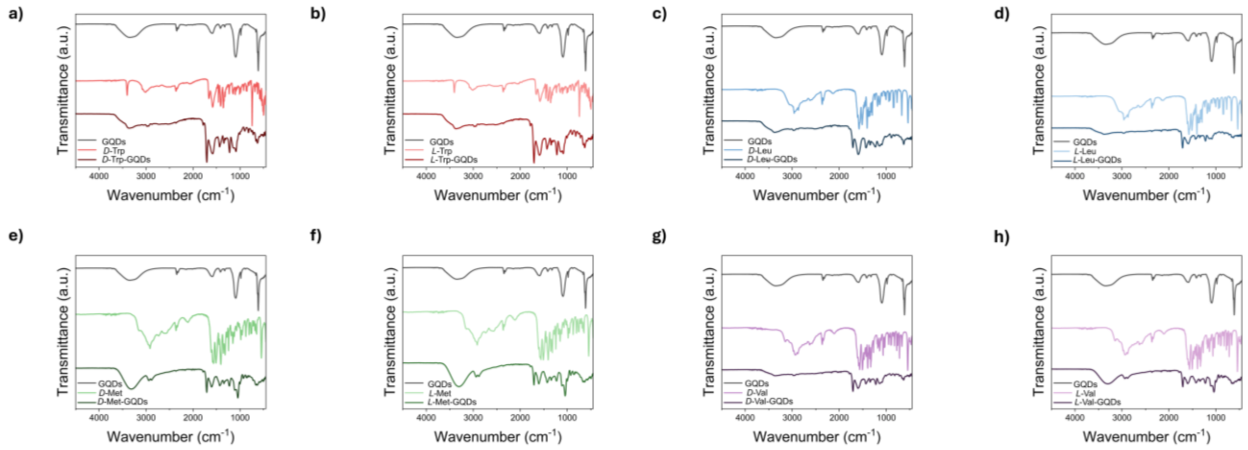


**Figure S3.** FTIR spectra of pristine GQDs and *D*- or *L*-amino acid–modified GQDs. a) *D*-Trp–GQDs, b) *L*-Trp–GQDs, c) *D*-Leu–GQDs, d) *L*-Leu–GQDs, e) *D*-Met–GQDs, f) *L*-Met–GQDs, g) *D*-Val–GQDs, and h) *L*-Val–GQDs. Characteristic amide vibrations at ~1730, 1545, and 1250 $cm^{-1}$ and C–H stretching bands at 2900–3050 $cm^{-1}$ confirm covalent conjugation of amino acids to GQD edges.

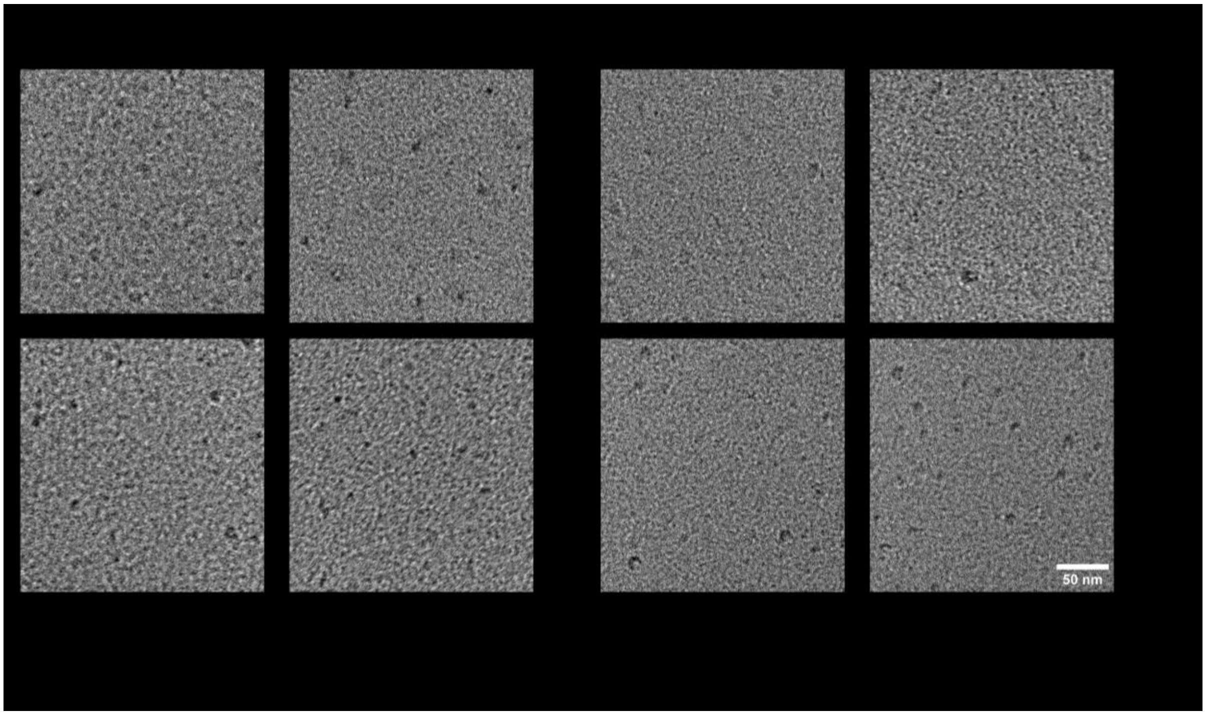


**Figure S4.** Representative Transmission electron microscopy (TEM) images of pristine and amino acid–modified GQDs( His, Cys, Ser and Thr). Images show uniform lateral dimensions (~6 nm) across variants, with no evidence of aggregation or major disruption of the graphene lattice. These results confirm that amino acid conjugation preserves the crystalline core while altering surface chemistry. Scale bars = 50 nm.

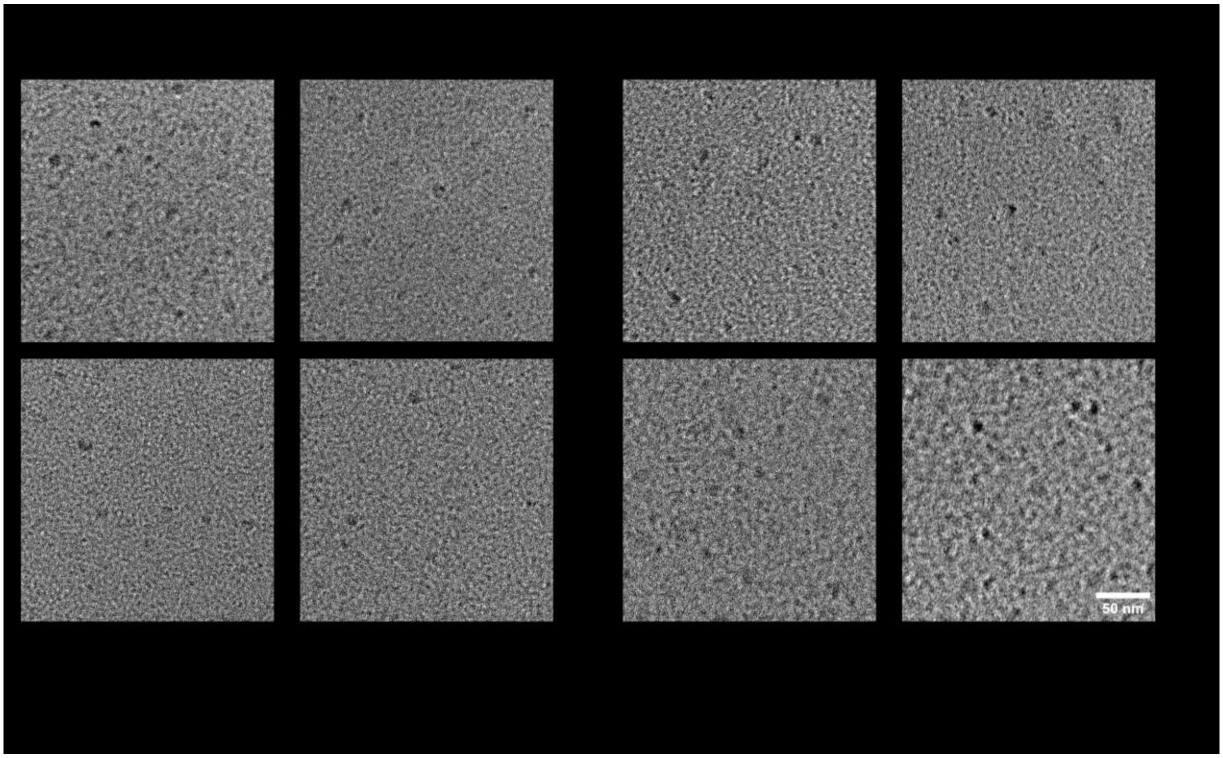


**Figure S5.** TEM images of amino acid–modified GQDs (Trp, Met, Leu and Val). All samples exhibit uniformly dispersed nanosheets with average lateral dimensions of ~6 nm, comparable to pristine GQDs. The absence of aggregation or lattice disruption indicates that covalent conjugation preserves the crystalline graphene while modifying edge chemistry. Scale bars = 50 nm.

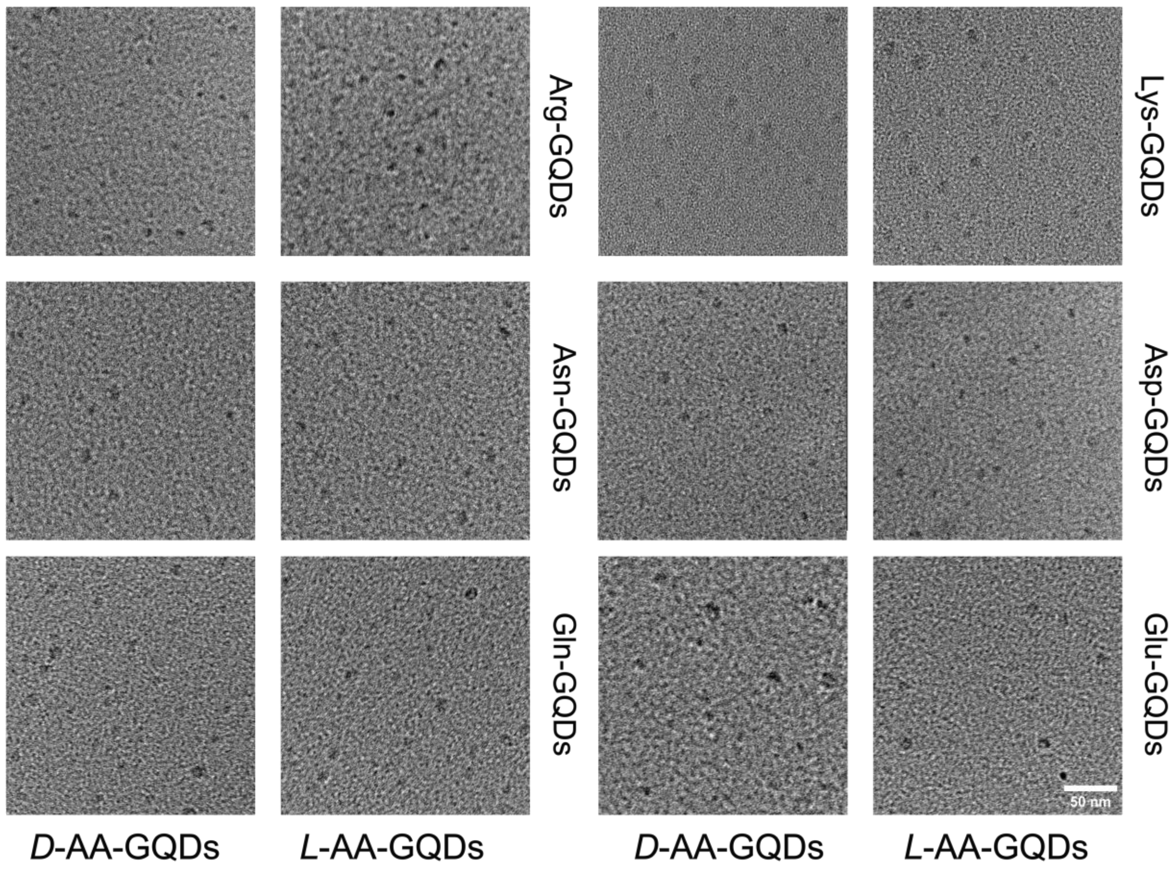


**Figure S6.** TEM images of chiral ligand functionalized GQDs (Arg, Lys, Asn, Asp, Gln, Glu). Samples show nanosheet structures with average lateral dimensions of ~6 nm, similar to pristine GQDs. No aggregation or lattice collapse is observed, confirming that edge functionalization preserves the crystalline graphene framework while modifying surface chemistry. Scale bars = 50 nm.

**Table S1.** Atomic force microscopy (AFM)-derived height distributions of pristine and chiral ligand functionalized GQDs. Values are reported as mean ± standard deviation (nm). Variants are grouped according to structural classification: hydrophobic (Met, Val, Leu), hydrophobic-special (Trp), twisted (Cys, Ser, Thr), saddle (His), positively charged (Arg, Lys), and polar/negatively charged (Asp, Glu, Gln, Asn). Height increases observed in Cys-, Ser-, Trp-, and His–GQDs indicate out-of-plane distortions associated with their respective chiral nanostructures.

| Name | Height (nm) | Nanostructure Type |
|---|---|---|
| **GQDs** | 1.17 ± 0.29 | N/A |
| ***D*-Met-GQDs** | 1.18 ± 0.10 | Unblocked |
| ***L*-Met-GQDs** | 1.21 ± 0.13 | Unblocked |
| ***D*-Val-GQDs** | 1.11 ± 0.34 | Unblocked |
| ***L*-Val-GQDs** | 1.16 ± 0.23 | Unblocked |
| ***D*-Leu-GQDs** | 1.23± 0.07 | Unblocked |
| ***L*-Leu-GQDs** | 1.22 ± 0.33 | Unblocked |
| ***D*-Trp-GQDs** | 1.56 ± 0.28 | Hybrid |
| ***L*-Trp-GQDs** | 1.74 ± 0.49 | Hybrid |
| ***D*-Cys-GQDs** | 1.58 ± 0.20 | Twisted |
| ***L*-Cys-GQDs** | 1.42 ± 0.19 | Twisted |
| ***D*-Ser-GQDs** | 1.62 ± 0.51 | Twisted |
| ***L*-Ser-GQDs** | 1.56 ± 0.60 | Twisted |
| ***D*-Thr-GQDs** | 1.12 ± 0.10 | Twisted |
| ***L*-Thr-GQDs** | 1.24 ± 0.17 | Twisted |
| ***D*-His-GQDs** | 1.94 ± 0.33 | Saddle |
| ***L*-His-GQDs** | 2.04 ± 0.74 | Saddle |
| ***D*-Arg-GQDs** | 1.17 ± 0.26 | Random |
| ***L*-Arg-GQDs** | 1.16 ± 0.26 | Random |
| ***D*-Lys-GQDs** | 1.16 ± 0.11 | Random |
| ***L*-Lys-GQDs** | 1.22 ± 0.23 | Random |
| ***D*-Asp-GQDs** | 1.03 ± 0.22 | Twisted Boat |
| ***L*-Asp-GQDs** | 1.11 ± 0.14 | Twisted Boat |
| ***D*-Glu-GQDs** | 1.24 ± 0.11 | Twisted Boat |
| ***L*-Glu-GQDs** | 1.23 ± 0.11 | Twisted Boat |
| ***D*-Gln-GQDs** | 1.16± 0.14 | Twisted Boat |
| ***L*-Gln-GQDs** | 1.13± 0.09 | Twisted Boat |
| ***D*-Asn-GQDs** | 1.23 ± 0.14 | Twisted Boat |
| ***L*-Asn-GQDs** | 1.11 ± 0.09 | Twisted Boat |

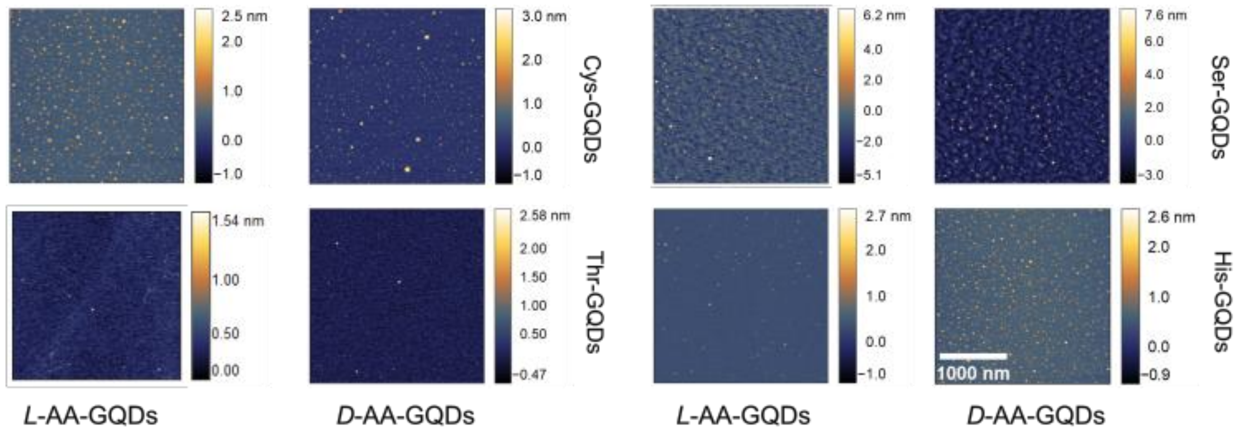


**Figure S7.** Representative AFM images of amino acid–modified GQDs. Cys-, Thr-, and Ser-GQDs exhibited twisted conformations, while His-GQDs formed saddle-shaped nanostructures. In both cases, increased heights relative to pristine GQDs confirmed localized out-of-plane distortions associated with emergent chirality.

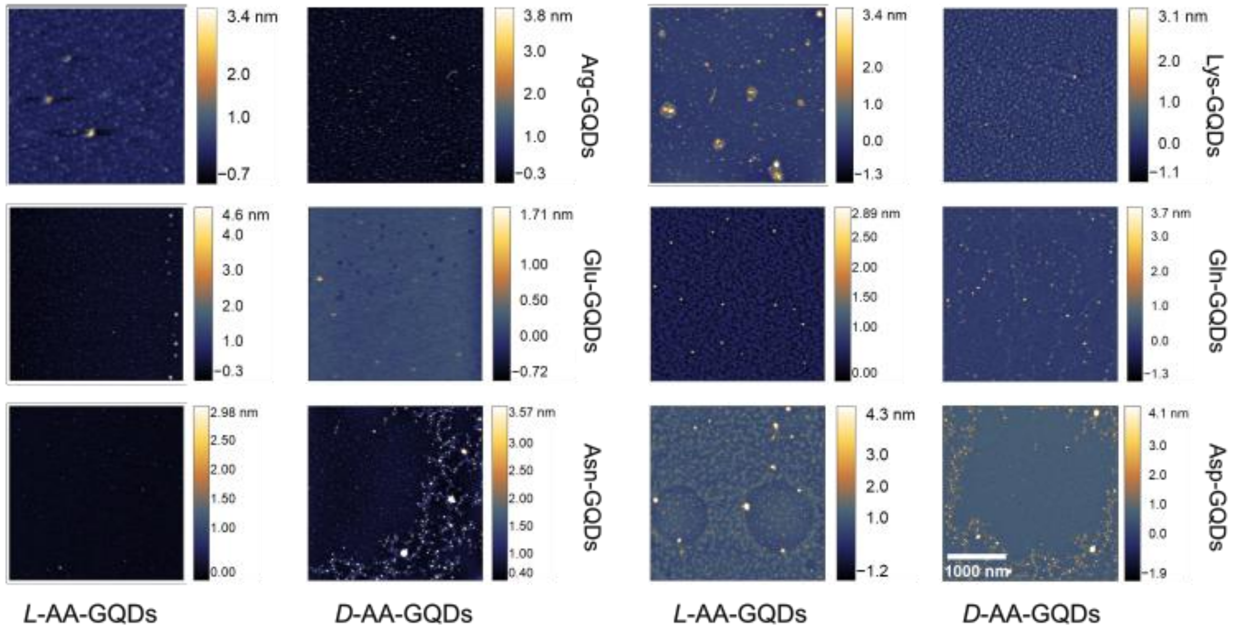


**Figure S8.** Representative AFM images of amino acid–modified GQDs. Lys- and Arg-GQDs (Random Structure), as well as Asp-, Asn-, Gln-, and Glu-GQDs (twisted Boat structure), showed no detectable out-of-plane distortions, indicating the absence of stable chiral nanostructures in these variants.

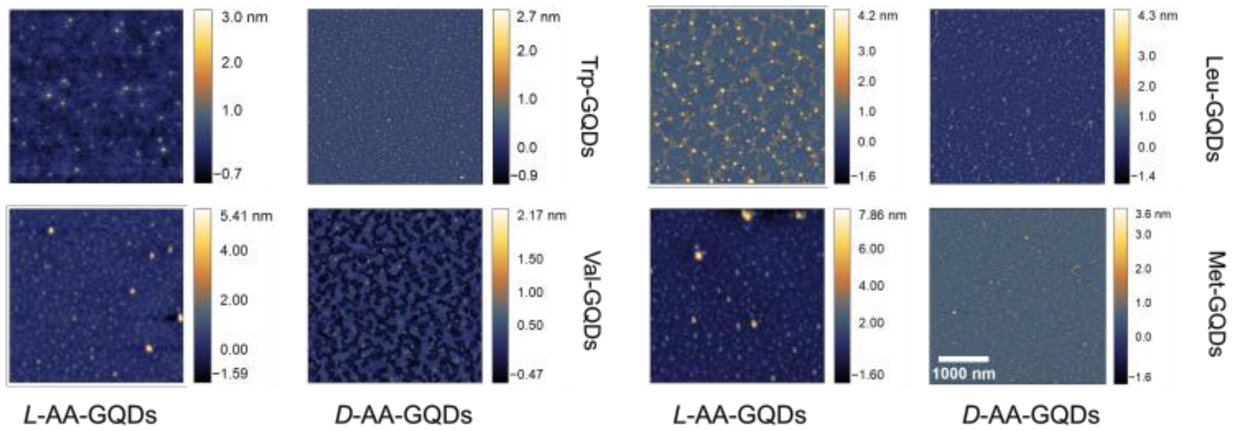


**Figure S9.** Representative AFM images of amino acid–modified GQDs. Trp-GQDs exhibited hybrid conformations with detectable out-of-plane distortions, whereas Val-, Leu-, and Met-GQDs showed no measurable distortions and remained unbuckled, indicating the absence of stable chiral nanostructures in these variants.

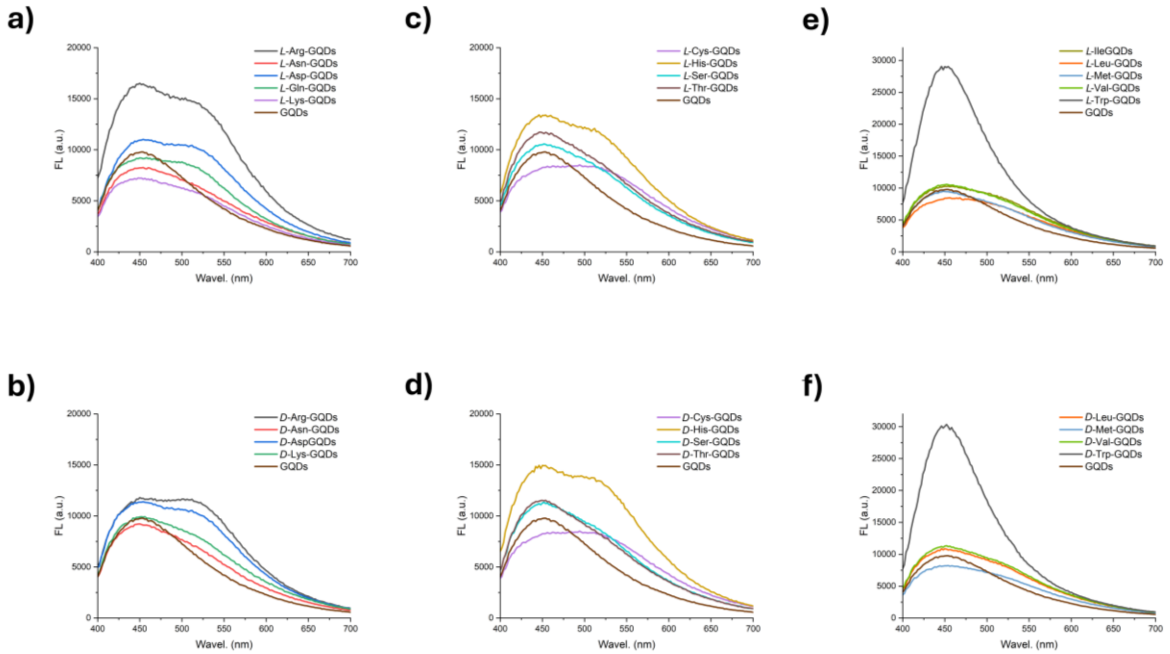


**Figure S10.** Photoluminescence (PL) spectra of pristine and amino acid–modified GQDs excited at 360 nm. a) *L*-Arg–, *L*-Asn–, *L*-Asp–, *L*-Gln- and *L*-Lys–GQDs. b) *D*-Arg–, *D*-Asn–, *D*-Asp–, *D*-Gln- and *D*-Lys–GQDs. c) L-Cys–, *L*-His–, *L*-Ser–, and *L*-Thr–GQDs. d) *D*-Cys–, *D*-His–, *D*-Ser–, and *D*-Thr–GQDs. e) *L*-Ile–, *L*-Leu–, *L*-Met–, *L*-Trp-, and *L*-Val–GQDs. f) *D*-Leu–, *D*-Met–, *D*-Val–, and *D*-Trp–GQDs. All modified GQDs exhibit re*D*-shifted emission relative to pristine GQDs, with emission maxima in the 450–550 nm range, consistent with edge functionalization and altered electronic environments.

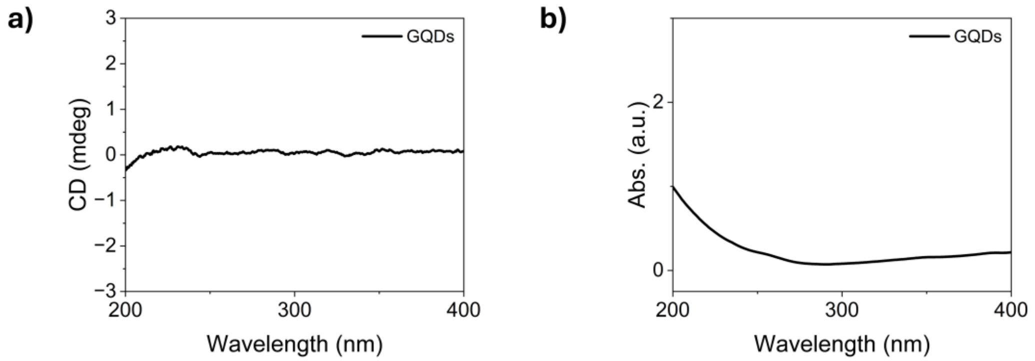


**Figure S11.** a) Circular dichroism (CD) spectrum and b) UV–Vis absorbance spectrum of pristine GQDs. The CD spectrum shows no detectable chiroptical activity within 200–400 nm, confirming the achiral nature of the unmodified GQDs.

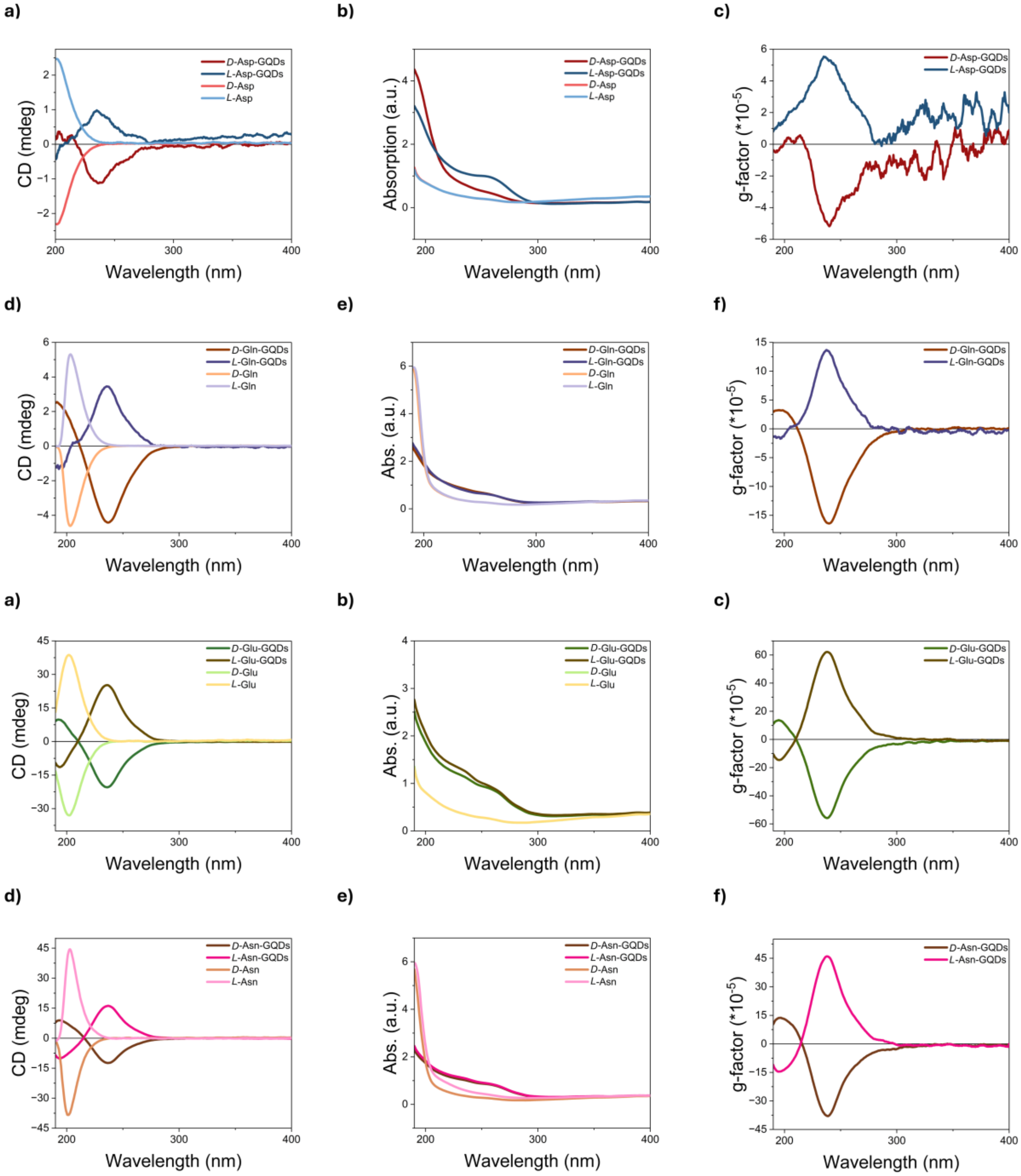


**Figure S12.** Circular dichroism (CD), UV–vis absorption, and anisotropy (g-factor) spectra of amino acid–modified GQDs compared with pristine GQDs. a–c) Asp–GQDs, d–f) Gln–GQDs, g–i) Glu–GQDs, j–l) Asn–GQDs. Each set shows CD response (left), absorption spectra (middle), and calculated g-factor (right), Data highlights characteristic chiroptical responses across twisted boat and random nanostructures, confirming ligand-dependent optical activity.

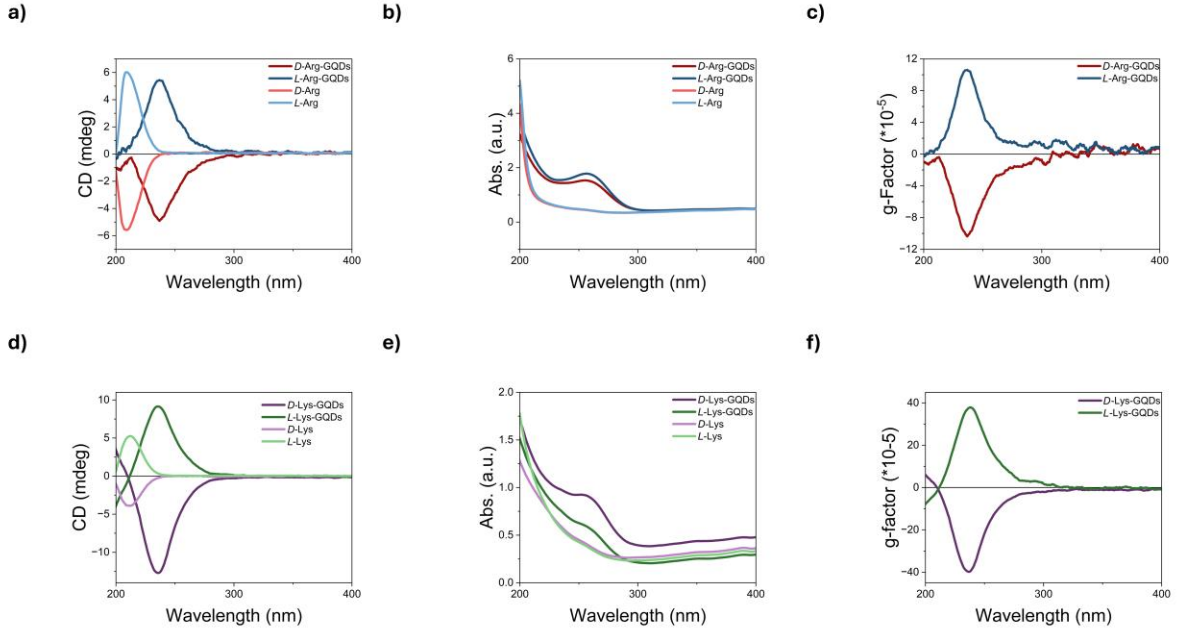


**Figure S13.** Circular dichroism (CD), UV–vis absorption, and anisotropy (g-factor) spectra of amino acid–modified GQDs compared with pristine GQDs. a–c) Arg–GQDs, d–f) Lys–GQDs. Each set shows CD response (left), absorption spectra (middle), and calculated g-factor (right), Data highlights characteristic chiroptical responses across twisted boat and random nanostructures, confirming ligand-dependent optical activity.

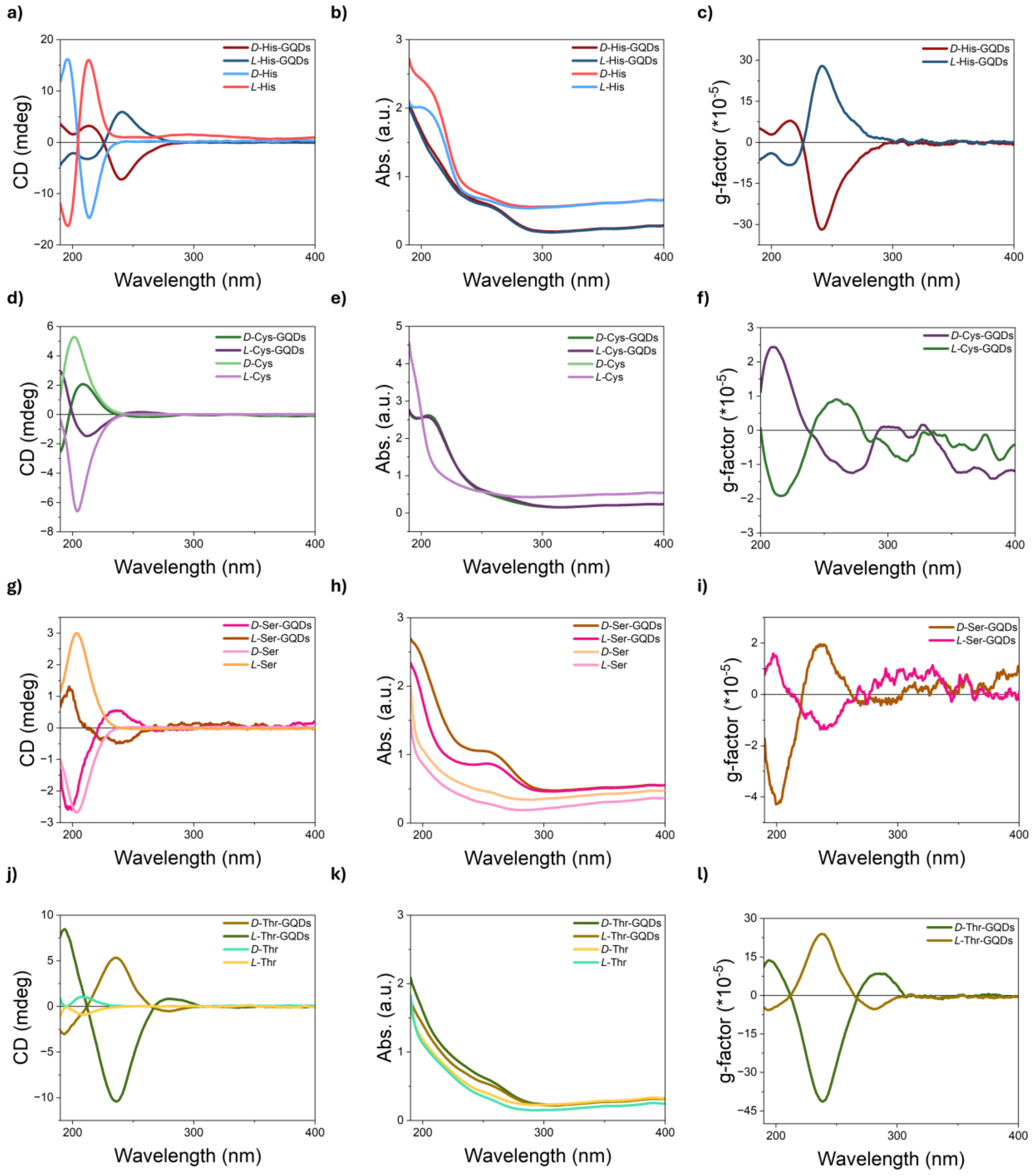


**Figure S14.** CD, UV–vis absorption, and anisotropy (g-factor) spectra of amino acid–modified GQDs. a–c) His–GQDs, d–f) Cys–GQDs, g–i) Ser–GQDs, j–l) Thr–GQDs Each panel set presents CD spectra (left), absorption profiles (middle), and corresponding g-factor plots (right). Data highlights characteristic chiroptical responses across twisted and saddle nanostructures, confirming ligand-dependent optical activity.

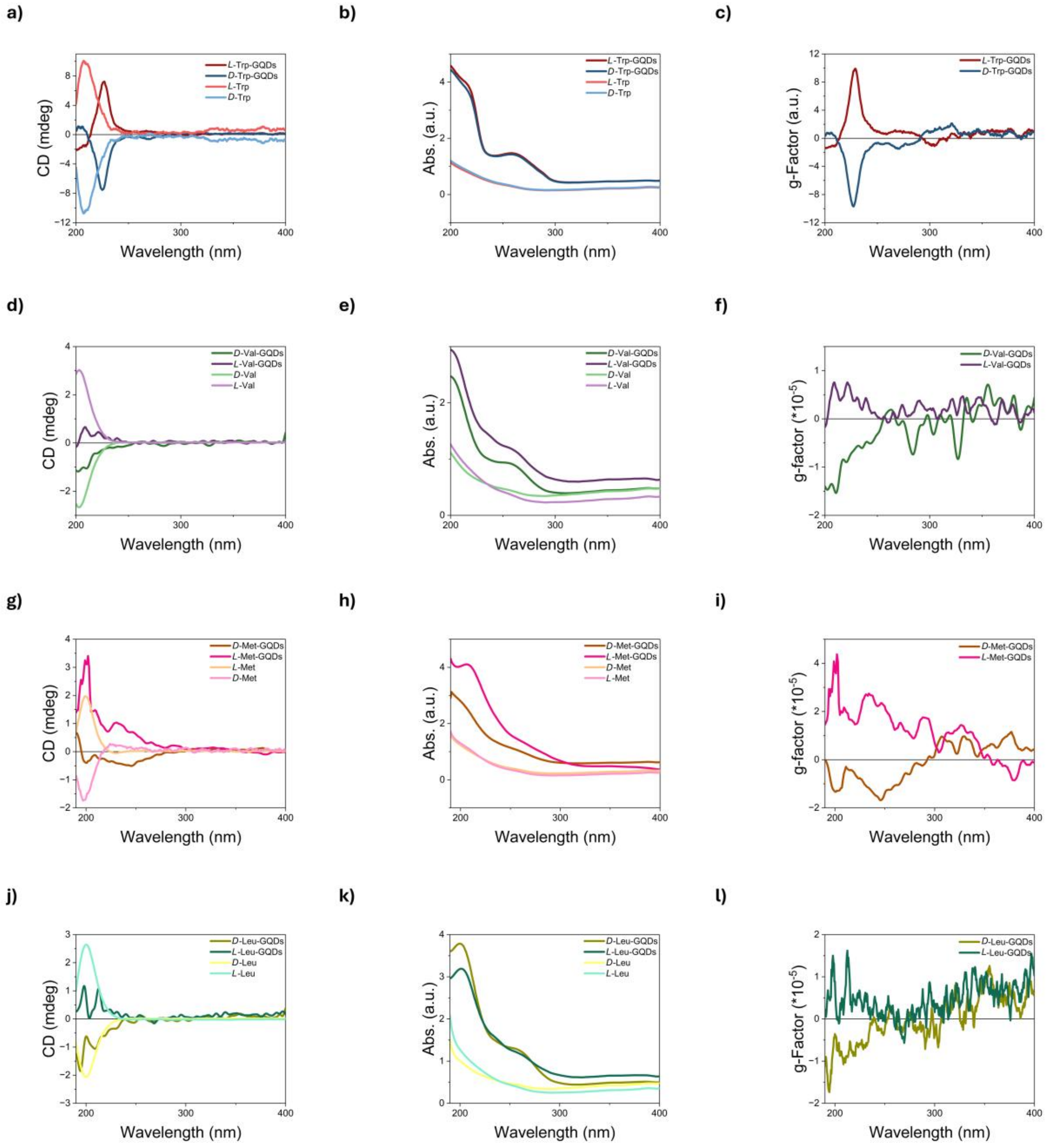


**Figure S15.** CD, UV–vis absorption, and anisotropy (g-factor) spectra of amino acid–modified GQDs. a–c) Trp–GQDs, d–f) Val–GQDs, g–i) Met–GQDs, and j–l) Leu–GQDs. Each panel set presents CD spectra (left), absorption profiles (middle), and g-factor plots (right). Data illustrate characteristic optical responses of hybrid and unbuckled systems, with Trp–GQDs showing transient structural contributions and hydrophobic residues (Val, Met, Leu) displaying weak, nonpersistent chiroptical signals.

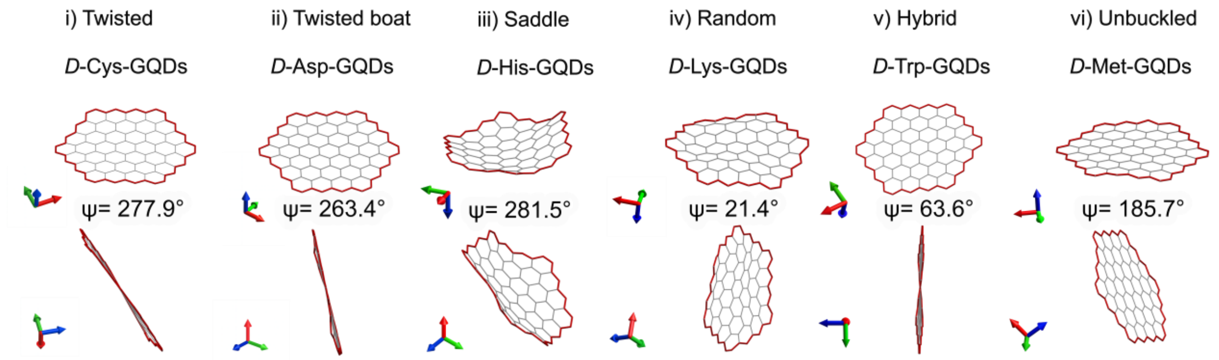


**Figure S16.** DFT-optimized geometries of *D*-amino-acid-modified GQDs showing distinct chiral distortions. The red outlines highlight the deformation of the graphene lattice upon amino acid conjugation. The calculated dihedral angle (ψ), Distinct nanostructural forms are observed: *D*-His-GQDs adopt a saddle-shaped configuration, *D*-Cys-GQDs exhibit a twisted structure, *D*-Trp-GQDs form a hybrid morphology, *D*-Met-GQDs remain unbuckled, *D*-Lys-GQDs display random ripples, and *D*-Asp-GQDs show a twisted-boat conformation.

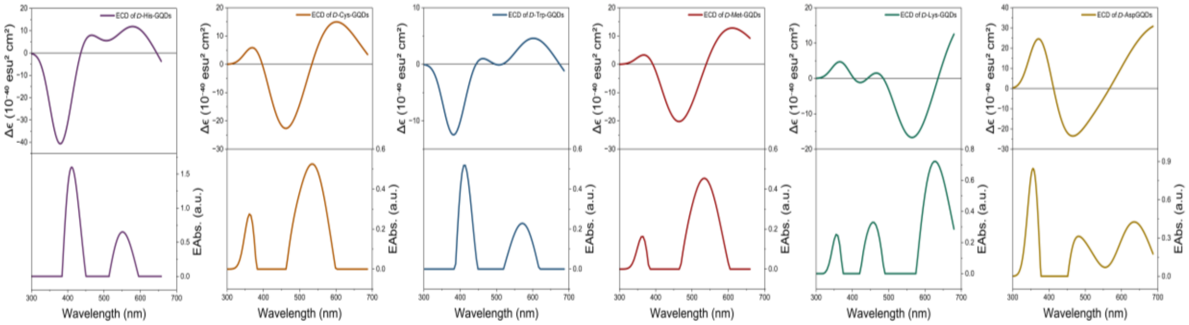


**Figure S17.** Time-dependent DFT (TD-DFT) simulated circular dichroism (top) and absorption (bottom) spectra of *D*-amino-acid–modified GQDs, calculated from the optimized ground-state geometries. The TD-DFT simulations reproduce the key spectral envelopes and relative intensities observed experimentally, confirming that the optimized geometries capture the principal chiroptical and electronic transitions of *D*-His-, *D*-Asp-, *D*-Lys-, *D*-Met-, *D*-Cys-, and *D*-Trp-GQDs. A systematic red shift of the experimental spectra relative to the computed ones is evident, consistent with the well-known underestimation of excitation energies by the B3LYP functional and similar hybrid methods for extended π-conjugated systems. Comparable offsets have been reported in other TD-DFT studies of carbon nanodots and graphene derivatives, supporting the reliability of the calculated trends.

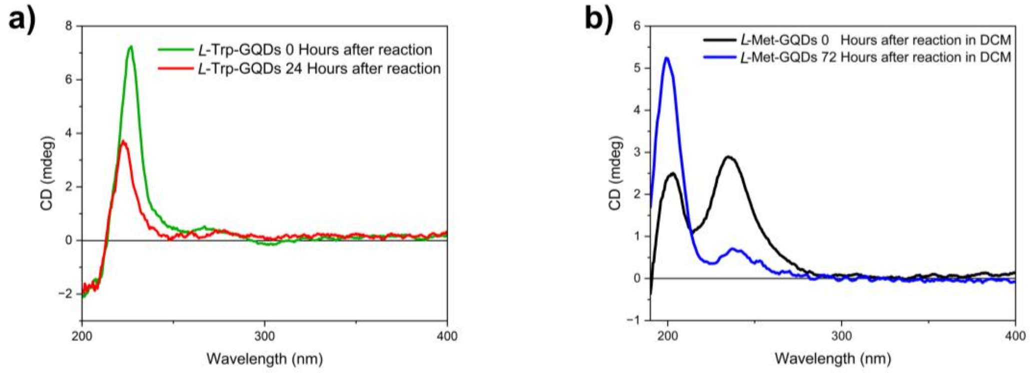


**Figure S18.** CD spectra of *L*-Met-GQDs synthesized in DCM and transferred into water, showing the gradual disappearance and folding of hydrophobic side chains onto the GQD surface over time. CD spectra of *L*-Trp-GQDs synthesized in DI water, showing the gradual disappearance and folding of hydrophobic side chains onto the GQD surface over time.

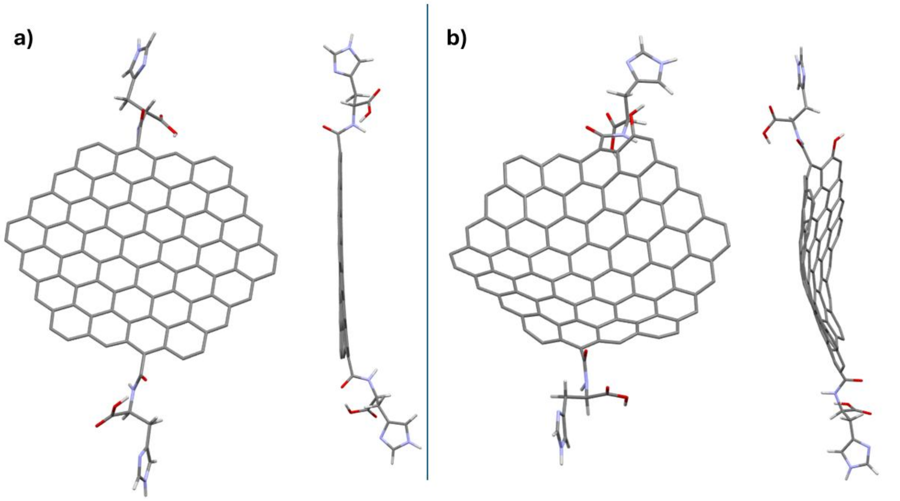


**Figure S19.** DFT-optimized geometries of His–GQDs. a) Racemic DL-His–GQDs showing no out-of-plane distortion and lacking nanostructural chirality. b) *D*-His–GQDs exhibiting a saddle-shaped geometry with pronounced vertical deformation, representing a stable chiral nanostructure.

**Table S2.** Total Puckering amplitude of different types of chiral GQDs based on the DFT optimized geometry.

| Name | Total Puckering (Q) | Nanostructure Type |
|---|---|---|
| ***D*-Cys-GQDs** | 0.57 | Twisted |
| ***D*-Ser-GQDs** | 0.85 | Twisted |
| ***D*-Thr-GQDs** | 0.41 | Twisted |
| ***D*-His-GQDs** | 1.25 | Saddle |
| ***D*-Asp-GQDs** | 0.48 | Twisted Boat |
| ***D*-Glu-GQDs** | 0.61 | Twisted Boat |
| ***D*-Gln-GQDs** | 0.46 | Twisted Boat |
| ***D*-Asn-GQDs** | 0.43 | Twisted Boat |

**Table S3.** Summary of DFT-derived structural parameters for selected *D*-amino acid–modified GQDs. Handedness, puckering strength, and structural form were determined from optimized geometries. The total puckering amplitude (Q) quantifies deviation from planarity, while the phase angle (φ) defines handedness. Data highlights the diversity of structural motifs, ranging from twisted and saddle-shaped geometries to nearly planar forms.

| GQD | Handedness | Puckering Strength | Structural Form | $Q$ | $\varphi$ (°) |
|---|---|---|---|---|---|
| *D*-Lys-GQDs | Right | Strong | Random wrinkled sheet | 1.52 | 90 |
| *D*-His-GQDs | Left | Strong | Saddle or helical concavity | 1.4 | -62 |
| *D*-Asp-GQDs | Left | Mild | Twisted boat structure | 0.48 | -71 |
| *D*-Cys-GQDs | Left | Mild | Low curvature twist | 0.59 | -80 |
| *D*-Trp-GQDs | Left | Mild | Nearly planar and chair structure | 0.34 | -56 |
| *D*-Met-GQDs | Right | Weak | Nearly planar and chair structure | 0.2 | 90 |

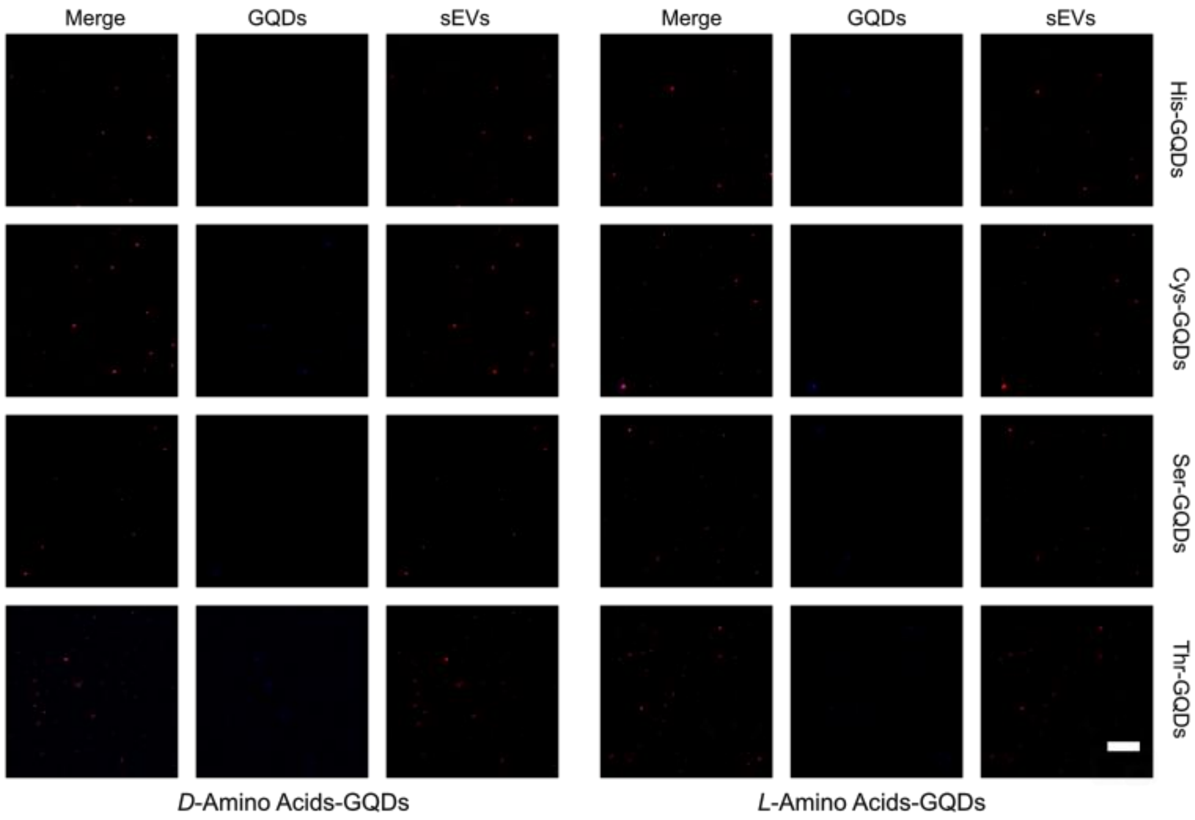


**Figure S20.** Confocal fluorescence microscopy images of small extracellular vesicles (sEVs) incubated with ligand–functionalized GQDs forming saddle-shaped (His-GQDs) and twisted (Cys-, Ser-, Thr-GQDs) nanostructures. *L*- and *D*-amino acids generated stable chiral nanostructures, with *D*-forms producing left-handed conformations that exhibited enhanced permeation into sEVs compared to the corresponding *L*-forms. Scale bar: 10 μm.

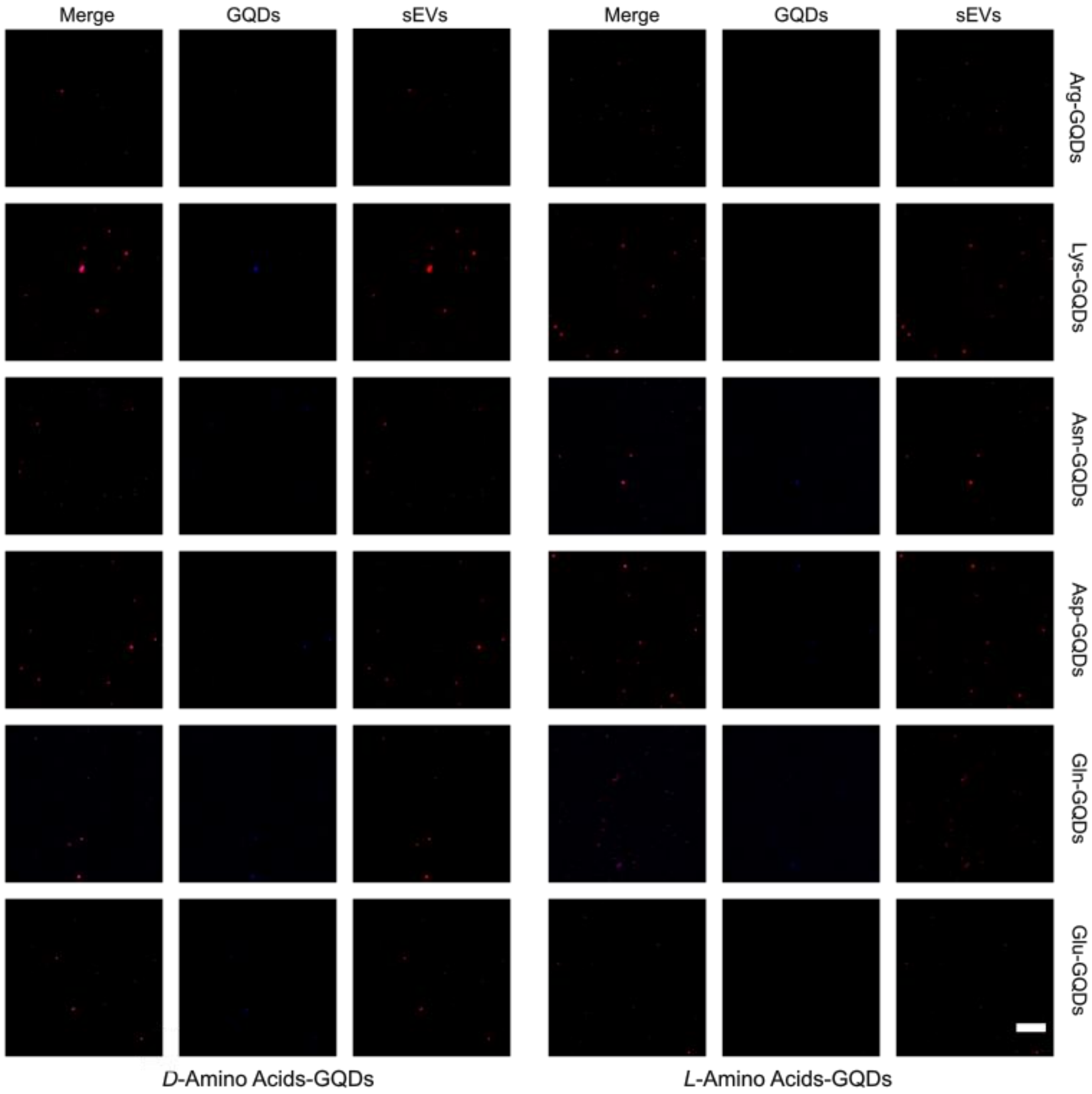


**Figure S21.** Confocal fluorescence microscopy images of sEVs incubated with amino acid–modified GQDs. Random-structure GQDs (Lys-, Arg-) showed minimal vesicle association with no difference between *L*- and *D*-forms. In contrast, twisted-boat GQDs (Asp-, Asn-, Gln-, Glu-) exhibited stronger permeation in the *D*-form, consistent with the formation of left-handed chiral nanostructures that enhance sEV entry. Scale bar: 10 μm.

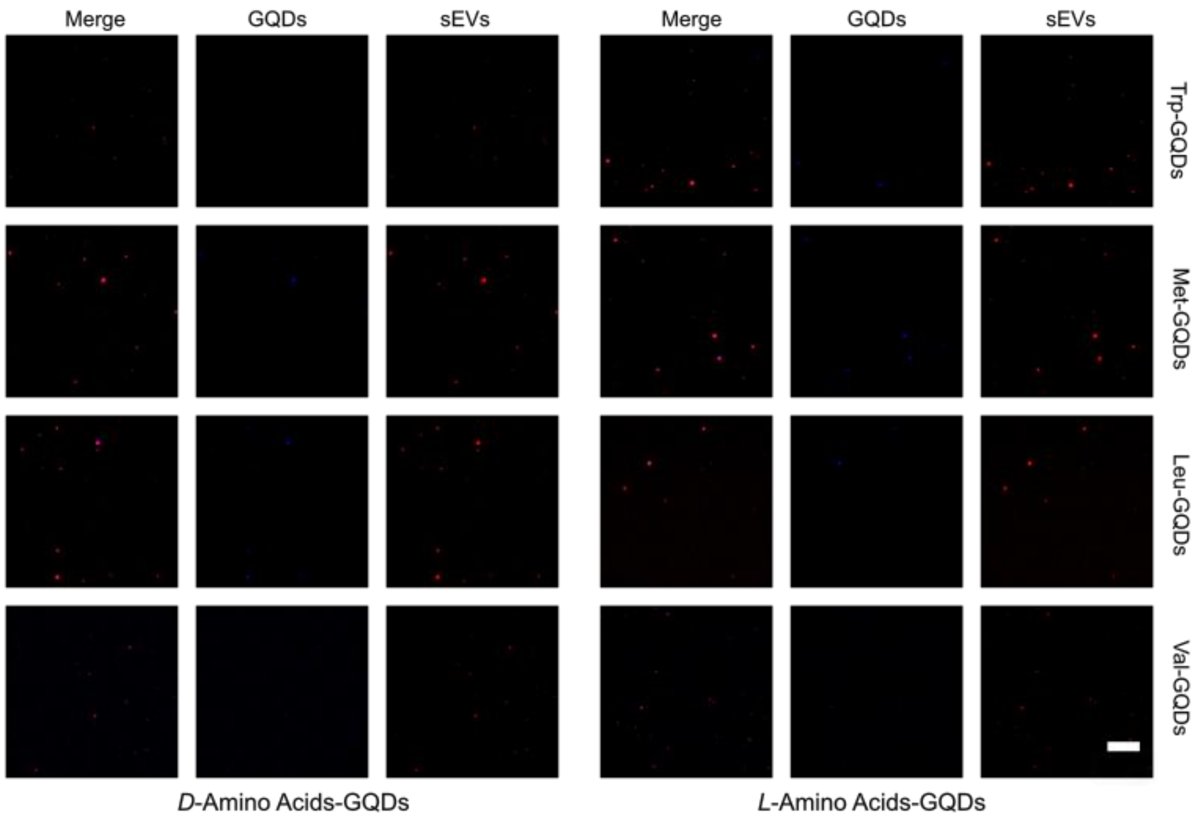


**Figure S22.** Confocal fluorescence microscopy images of sEVs incubated with hydrophobic amino acid–modified GQDs (Trp-, Met-, Leu-, and Val-GQDs). These variants form achiral or unbuckled nanostructures, and no significant differences in sEV permeation were observed between *L*- and *D*-enantiomers. Scale bar: 10 μm.

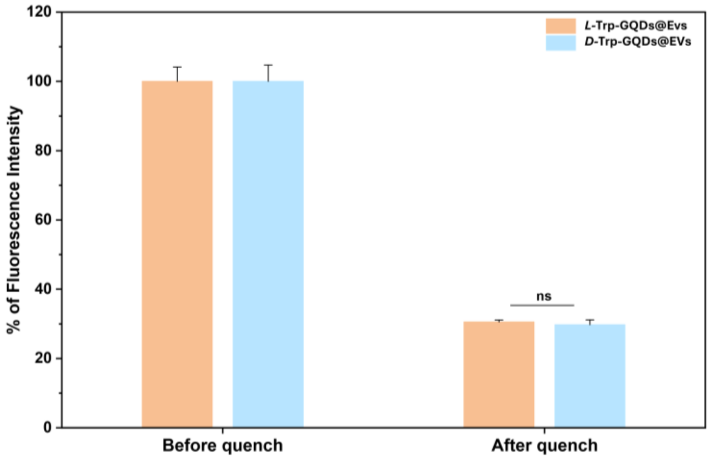


**Figure S23**. Fluorescence intensity of chiral Trp-GQDs before and after trypan blue quenching.

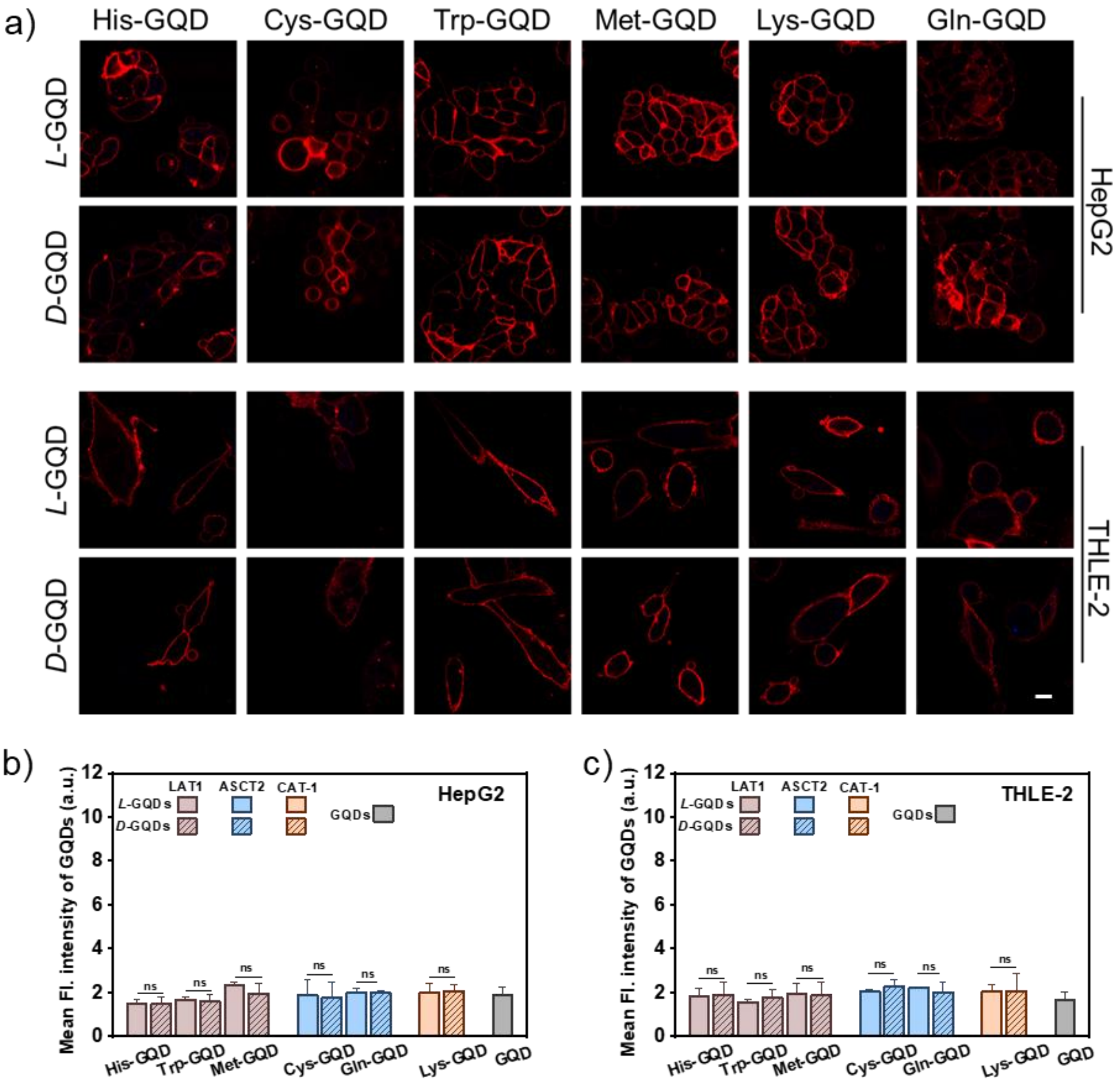


**Figure S24.** Cellular uptake of amino-acid–conjugated GQDs at 4 ℃. a) Representative confocal images of HepG2 and THLE-2 cells after 4 h incubation with the indicated chiral GQD variants (5 μM). DiD labels the plasma membrane. Scale bars: 10 μm. (*n* = 5 ). Quantification of cellular GQD fluorescence for b)HepG2 and c)THLE-2 corresponding to panel (a). Data are mean ± SD. Statistical differences were analyzed by a Student's two-sided t-test.
